# Public T-Cell Receptors (TCRs) Revisited by Analysis of the Magnitude of Identical and Highly-Similar TCRs in Virus-Specific T-Cell Repertoires of Healthy Individuals

**DOI:** 10.1101/2021.11.29.470325

**Authors:** Wesley Huisman, Lois Hageman, Didier A.T. Leboux, Alexandra Khmelevskaya, Grigory A. Efimov, Marthe C.J. Roex, Derk Amsen, J.H.F. Falkenburg, Inge Jedema

**Affiliations:** Department of Hematology, Leiden University Medical Center, The Netherlands; Department of Hematopoiesis, Sanquin Research and Landsteiner Laboratory for Blood Cell Research, Amsterdam, The Netherlands; Laboratory of Transplantation Immunology, National Research Center for Hematology, Moscow, Russia

**Keywords:** Virus-specific T cells, T-cell Receptors, TCRs, Public, Computational analysis, CDR3-region

## Abstract

Since multiple different T-cell receptor (TCR) sequences can bind to the same peptide-MHC combination and the number of TCR-sequences that can theoretically be generated even exceeds the number of T cells in a human body, the likelihood that many public identical (PUB-I) TCR-sequences frequently contribute to immune responses has been estimated to be low. Here, we quantitatively analyzed the TCR-repertoires of 190 purified virus-specific memory T-cell populations, directed against 21 antigens of Cytomegalovirus, Epstein-Barr virus and Adenovirus isolated from 29 healthy individuals, and determined the magnitude, defined as prevalence within the population and frequencies within individuals, of PUB-I TCR and of TCR-sequences that are highly-similar (PUB-HS) to these PUB-I TCR-sequences. We found that almost one third of all TCR nucleotide-sequences represented PUB-I TCR amino-acid (AA) sequences and found an additional 12% of PUB-HS TCRs differing by maximally 3 AAs. We illustrate that these PUB-I and PUB-HS TCRs were structurally related and contained shared core-sequences in their TCR-sequences. We found a prevalence of PUB-I and PUB-HS TCRs of up to 50% among individuals and showed frequencies of virus-specific PUB-I and PUB-HS TCRs making up more than 10% of each virus-specific T-cell population. These findings were confirmed by using an independent TCR-database of virus-specific TCRs. We therefore conclude that the magnitude of the contribution of PUB-I and PUB-HS TCRs to these virus-specific T-cell responses is high. Because the T cells from these virus-specific memory TCR-repertoires were the result of successful control of the virus in these healthy individuals, these PUB-HS TCRs and PUB-I TCRs may be attractive candidates for immunotherapy in immunocompromised patients that lack virus-specific T cells to control viral reactivation.

**Significance statement:** Public T-cell responses, in which T cells expressing the same T-cell receptor (TCR) are found in different individuals, have been described. However, the magnitude of the contribution of these TCRs to immune responses, defined as prevalence within the population and frequencies within individuals, is not known. In this study we characterized and quantified public T-cell responses within virus-specific memory T cells of healthy individuals by determining identical and highly-similar TCRs recognizing the same antigen and sharing conserved CDR3 motifs. The magnitude of public T-cell responses was surprisingly high and we argue that these dominant TCRs with shared core-sequences could be utilized for diagnostic purposes and may provide attractive TCRs to be used for immunotherapy in immunocompromised patients.

## Introduction

Human virus-specific CD8^pos^ T cells express heterodimeric alpha(α)/beta(β) TCRs that can specifically recognize viral peptides in the context of HLA-class-I molecules(1). The TCRα- and the TCRβ-chain repertoires are highly variable due to the genetic recombination process involved in their generation. For the TCRβ-chains, recombination of 1 of 48 functional T-cell Receptor Beta Variable (TRBV), 1 of 2 functional T-cell receptor Beta Diversity (TRBD) and 1 of 12 functional T-cell Receptor Beta Joining (TRBJ) gene segments leads to a V-D-J reading frame(2). The TCRα-chains are generated by a similar recombination process with the exception of a diversity gene, resulting in a V-J reading frame(3). Insertion of template-independent nucleotides between the recombined segments (junctional region) results in a significant further increase in variability(4). The sequence around these junctions encodes for the Complementary Determining Region 3 (CDR3), a loop that reaches out and interacts with a peptide embedded in an HLA molecule, together with the loops of the CDR1 and CDR2 regions, which are fixed within the germline variable gene sequence(5, 6). It has been calculated that these gene rearrangements could potentially generate a repertoire of 10^15^-10^20^ unique TCRs that may interact with all possible peptide-HLA complexes(7).

Pathogenic viruses like Cytomegalovirus (CMV), Epstein-Barr virus (EBV) and human Adenovirus (AdV) can infect individuals for life by staying latently present in target cells after a primary infection. To control these viruses, a population of virus-specific memory T cells has to develop from the vast naïve T-cell repertoire. Due to the high diversity of the naïve T-cell repertoire(8), T-cell responses against the many potential viral epitopes presented in multiple HLA alleles may be composed of a large variety of different TCRs. Indeed, when naïve UCB-derived T cells were stimulated *in vitro* to generate *de novo* responses against proteins from CMV or Human Immunodeficiency Virus (HIV), this resulted in responding virus-specific T-cell populations with a highly diverse repertoire of TCRs, recognizing many different CMV(9) or HIV-derived peptides(10). However, from *ex vivo* analyses in adults it became clear that *in vivo* the virus-specific memory T-cell populations are shaped during control and clearance of the infection and target only a limited number of viral-peptides, as was shown for T-cell populations specific for viruses like CMV(11), EBV(12), AdV(13), Influenza A(14) and also more recently SARS-Cov-2(15, 16). Nevertheless, the multiple viral-peptides that are targeted in the various HLA alleles make it theoretically unlikely that individuals would frequently share exactly the same virus-specific TCR, unless T cells expressing certain TCRs would favor control of infections and would therefore dominate the responses.

Evidence for selection of certain virus-specific TCR-expressing T cells in controlling viruses has come from several reports identifying identical TCR amino-acid (AA) sequences in dominant virus-specific memory T-cell populations in different individuals, designated as public TCR-sequences (from here on referred to as public-identical (PUB-I) TCR-sequences). These sequences have been found in CMV-specific T-cell responses(17–19), EBV-specific T-cell responses(20, 21) and SARS-Cov-2-specific T-cell responses(15, 22). In addition, some of these virus-specific T-cell populations also contained TCR AA-sequences that were highly-similar to the identical shared TCR AA-sequence (from here on referred to as highly-similar to PUB-I (PUB-HS) TCR-sequences). However, the magnitude, defined as prevalence within the population and frequencies within the individuals, of PUB-I and PUB-HS TCR-sequences is not known. The total combined PUB-I and PUB-HS TCR-sequences likely reflect the viral-antigen-specific T-cell repertoire that most optimally recognizes a specific peptide-HLA complex on infected target cells. Identification of such dominant TCRs with shared core-sequences could be utilized for diagnostic purposes or for the design of future immunotherapy purposes including TCR-gene transfer.

The aim of our study was to quantitively analyze the magnitude of PUB-I and PUB-HS TCR-sequences within the virus-specific TCR-repertoires of CMV, EBV and AdV-specific CD8^pos^ T cells. We confirmed that healthy individuals generate many different virus-specific TCRs, illustrated by the >3000 TCR nucleotide-sequences that were found *ex vivo* in virus-specific memory T-cell populations. However, a significant part of the virus-specific TCR-repertoires contained PUB-I and PUB-HS TCR nucleotide-sequences. For 19 out of the 21 different virus-specificities, PUB-I and PUB-HS TCRs could be found in multiple individuals. The AAs of these PUB-HS TCRs varied on specific positions in the CDR3β-region, while maintaining a conserved core-AA-sequence that was also present in the respective PUB-I TCR. We showed that the prevalence of PUB-I and PUB-HS TCRs among healthy individuals was more than 40% of the total virus-specific TCR-repertoire and that they occurred at a median frequency of 13.1% within each virus-specific T-cell population, which was unexpectedly high. We identified conserved TCR core-AA-sequences for each specificity that could be used for diagnostic purposes looking at anti-viral immune responses. Additionally, PUB-I or PUB-HS TCRs with the highest frequencies in healthy individuals might be utilized to develop off-the-shelf immunotherapeutics (using TCR-gene transfer) to effectively control CMV, EBV or AdV-infections or reactivations in immunocompromised patients.

## Results

### Generation and validation of a library of TCR-sequences derived from FACsorted virus-specific T-cell populations

To examine the composition of the virus-specific TCR-repertoires in different individuals, the CDR3β-regions of purified expanded pMHC-tetramer-binding virus-specific T-cell populations were sequenced **(Supplementary Figure 1).** We analyzed the TCR-repertoires of CMV, EBV and AdV-specific T cells, restricted to four prevalent HLA alleles (HLA-A*01:01, HLA-A*02:01, HLA-B*07:02 and HLA-B*08:01) and specific for 21 different peptides (**Table 1**). Purified CMV, EBV and AdV-specific T-cell populations targeting CMV (n=8), EBV (n=10) or AdV (n=3)-derived peptides were isolated from 17 HLA-A*01:01/B*08:01^pos^ individuals and 12 HLA-A*02:01/B*07:02^pos^ individuals **(Table 1 and Table 2).** In total, 190 virus-specific T-cell populations, each targeting a single viral antigen, were successfully isolated and showed high purity (>97% pMHC-tetramer positive). Sequencing of the CDR3β-regions of these virus-specific T-cell populations resulted in 3346 CDR3β nucleotide-sequences that occurred at frequencies of more than 0.1%. 135 of these nucleotide-sequences were present at high frequencies (>5%) within one specificity, but were also found at low frequencies (around 0.1%) in another specificity, indicating contamination due to FACSorting impurities. These low frequency nucleotide-sequences were discarded from further analysis. 41 nucleotide-sequences were present at low frequencies in two unrelated specificities and could not be correctly annotated and these 41 duplicates (n=82) were also discarded from the analysis. Therefore, a total of 3129 nucleotide-sequences could be annotated. In total, 2224 (71%) of these nucleotide-sequences represented unique CDR3β AA-sequences that were found in only one individual and 905 nucleotide-sequences (29%) resulted in 131 different PUB-I CDR3β AA-sequences that were found in two or more unrelated individuals **(Flowchart; Figure 1).**

**Table 1:**
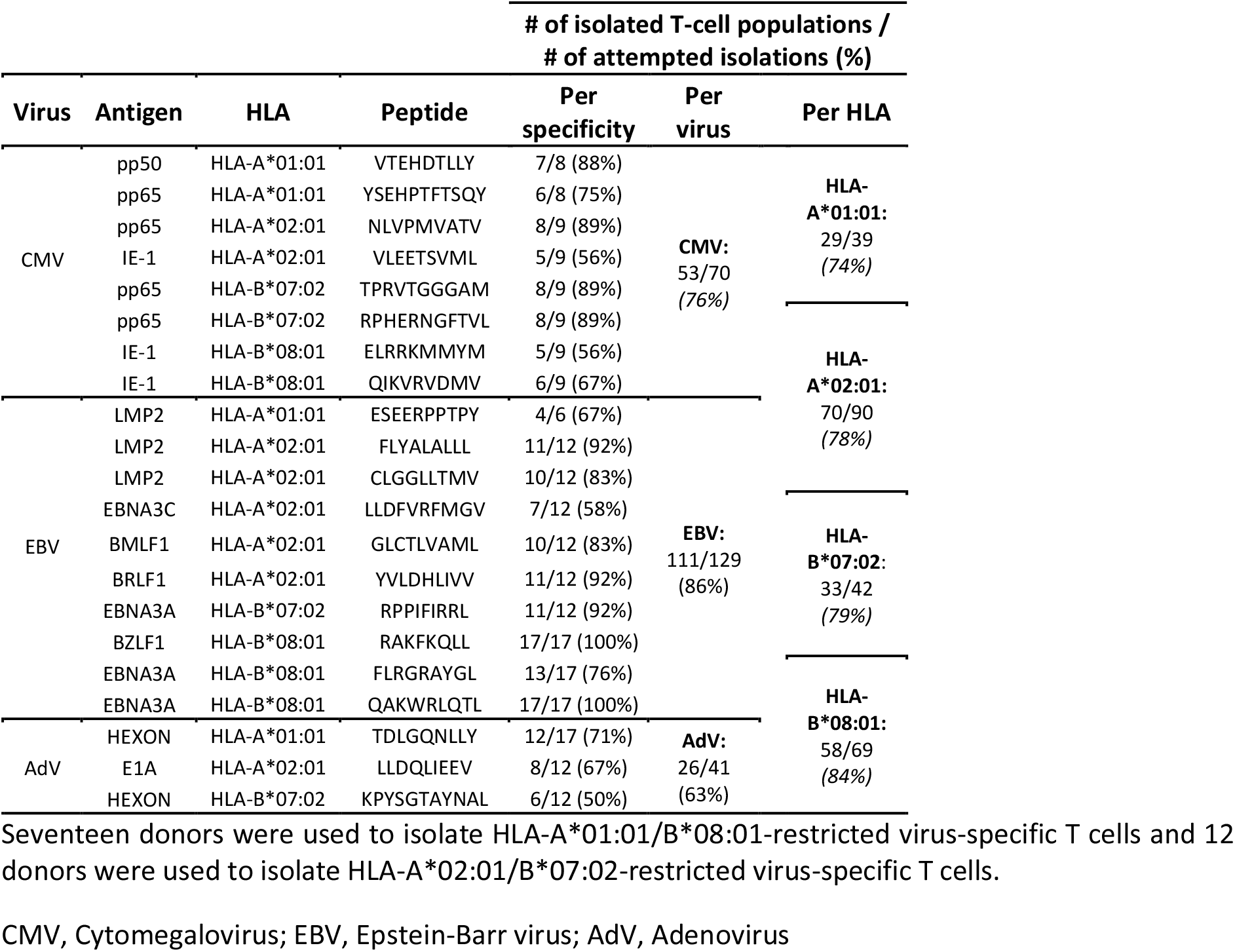
Number of isolated virus-specific T-cell populations.

**Table 2:** HLA typing and CMV/EBV-serostatus of healthy donors.

| # | CMV | EBV | HLA-A |  | HLA-B |  | HLA-C |  | HLA-DR |  | HLA-DQ |  | HLA-DP |  |
| --- | --- | --- | --- | --- | --- | --- | --- | --- | --- | --- | --- | --- | --- | --- |
| 1 | Pos | Pos | 01:01 | 30:01 | 08:01 | 13:02 | 07:01 | 06:02 | 03:01 | 07:01 | 02:01 | 02:02 | 04:01 | 09:01 |
| 2 | Pos | Pos | 01:01 |  | 08:01 |  | 07:01 |  | 03:01 | 01:02 | 02:01 | 05:01 | 01:01 | 04:01 |
| 3 | Pos | Pos | 01:01 |  | 08:01 |  | 07:01 |  | 03:01 | 15:01 | 02:01 | 06:02 | N.D |  |
| 4 | Pos | Pos | 01:01 |  | 08:01 |  | 07:01 |  | 03:01 |  | 02:01 |  | 04:01 |  |
| 5 | Pos | pos | 01:01 | 30:01 | 08:01 | 13:02 | 07:01 | 06:02 | 04:07 | 15:01 | 03:01 | 06:02 | 04:01 |  |
| 6 | Pos | Pos | 01:01 |  | 08:01 |  | 07:01 |  | 03:XX |  | 02:XX |  | N.D |  |
| 7 | Pos | pos | 01:01 | 30:01 | 08:01 | 13:02 | 07:01 | 06:02 | 03:01 | 04:01 | 02:01 | 03:01 | 04:01 |  |
| 8 | Pos | pos | 02:01 | 24:02 | 08:01 | 35:01 | 07:01 | 11:01 | 02:02 | 03:01 | 02:02 | 03:01 | 02:01 | 13:01 |
| 9 | Neg | Pos | 01:01 |  | 08:01 |  | 07:01 |  | 03:01 |  | 02:01 |  | 04:01 |  |
| 10 | Neg | Pos | 01:01 |  | 08:01 |  | 07:01 |  | 03:01 |  | 02:01 |  | 04:01 | 05:01 |
| 11 | Neg | Pos | 01:01 |  | 08:01 |  | 07:01 |  | 03:01 |  | 02:01 |  | 01:01 | 09:01 |
| 12 | Neg | Pos | 01:01 |  | 08:01 |  | 07:01 |  | 03:01 |  | 02:01 |  | 04:01 | 05:01 |
| 13 | Neg | Pos | 01:01 |  | 08:01 |  | 07:01 |  | 03:01 |  | 02:01 |  | 04:01 | 04:02/01 |
| 14 | Pos | Pos | 01:01 | 68:01 | 08:01 | 35:01 | 04:01 | 07:01 | 01:01 | 03:01 | 02 | 05:01 | 04:01 | 04:02 |
| 15 | Neg | Pos | 01:01 |  | 08:01 |  | 07:0 |  | 03:XX |  | 02:XX |  | N.D |  |
| 16 | Neg | pos | 01:01 | 24:02 | 08:01 | 35:01 | 04:01 | 07:01 | 03:01 | 08:01 | 02:01 | 04:02 | 04:01 |  |
| 17 | Neg | Pos | 01:01 |  | 08:01 |  | 07:01 |  | 03:XX |  | 02:XX |  | N.D |  |
| 18 | Pos | Pos | 02:01 |  | 07:02 | 44:02 | 07:02 | 05:01 | 15:01 | 04:01 | 06:02 | 03:01 | 04:XX | 02:01 |
| 19 | Pos | Pos | 02:01 |  | 07:02 |  | 07:02 |  | 15:01 |  | 06:02 |  | 04:01 |  |
| 20 | Pos | Pos | 02:01 | 03:01 | 07:02 |  | 07:02 |  | 15:01 |  | 06:02 |  | 04:01 | 03:01 |
| 21 | Pos | Pos | 02:01 |  | 07:02 |  | 07:02 |  | 15:01 |  | 06:02 |  | 04:01 | 05:01 |
| 22 | Pos | Pos | 02:01 |  | 07:02 |  | 07:02 |  | 15:01 |  | 06:02 |  | 02:01 | 04:01 |
| 23 | Pos | Pos | 02:01 |  | 07:02 |  | 07:02 |  | 15:01 |  | 06:02 |  | 04:01 |  |
| 24 | Pos | Pos | 02:01 | 03:01 | 07:02 | 44:02 | 07:02 | 05:01 | 15:01 | 01:01 | 06:02 | 05:01 | 04:01 | 14:01 |
| 25 | Pos | Pos | 02:01 |  | 07:02 |  | 07:02 |  | 07:01 | 15:01 | 03:03 | 06:02 | 04:01 | 13:01 |
| 26 | Pos | Pos | 02:01 |  | 07:02 |  | 07:02 |  | 15:XX |  | 06:XX |  | N.D |  |
| 27 | Neg | Pos | 02:01 |  | 07:02 |  | 07:02 |  | 15:01 |  | 06:02 |  | 04:01 |  |
| 28 | Neg | Pos | 02:01 |  | 07:02 |  | 07:02 |  | 15:01 |  | 06:02 |  | 04:01 | 13:01 |
| 29 | Neg | Pos | 02:01 |  | 07:02 |  | 07:02 |  | 15:XX |  | 06:XX |  | N.D |  |

Virus-specific T cells restricted to HLA-A\*01:01/HLA-B\*08:01 or HLA-A\*02:01/HLA-B\*07:02 were isolated from donors #1-17 and donors #18-29, respectively. CMV and EBV serostatus are indicated for each donor. HLA typing was determined either by serology, where the second digits could not be determined (XX), or with high resolution HLA typing unless indicated by N.D.. Blanks indicate homozygosity for the given allele.

**Figure 1:**
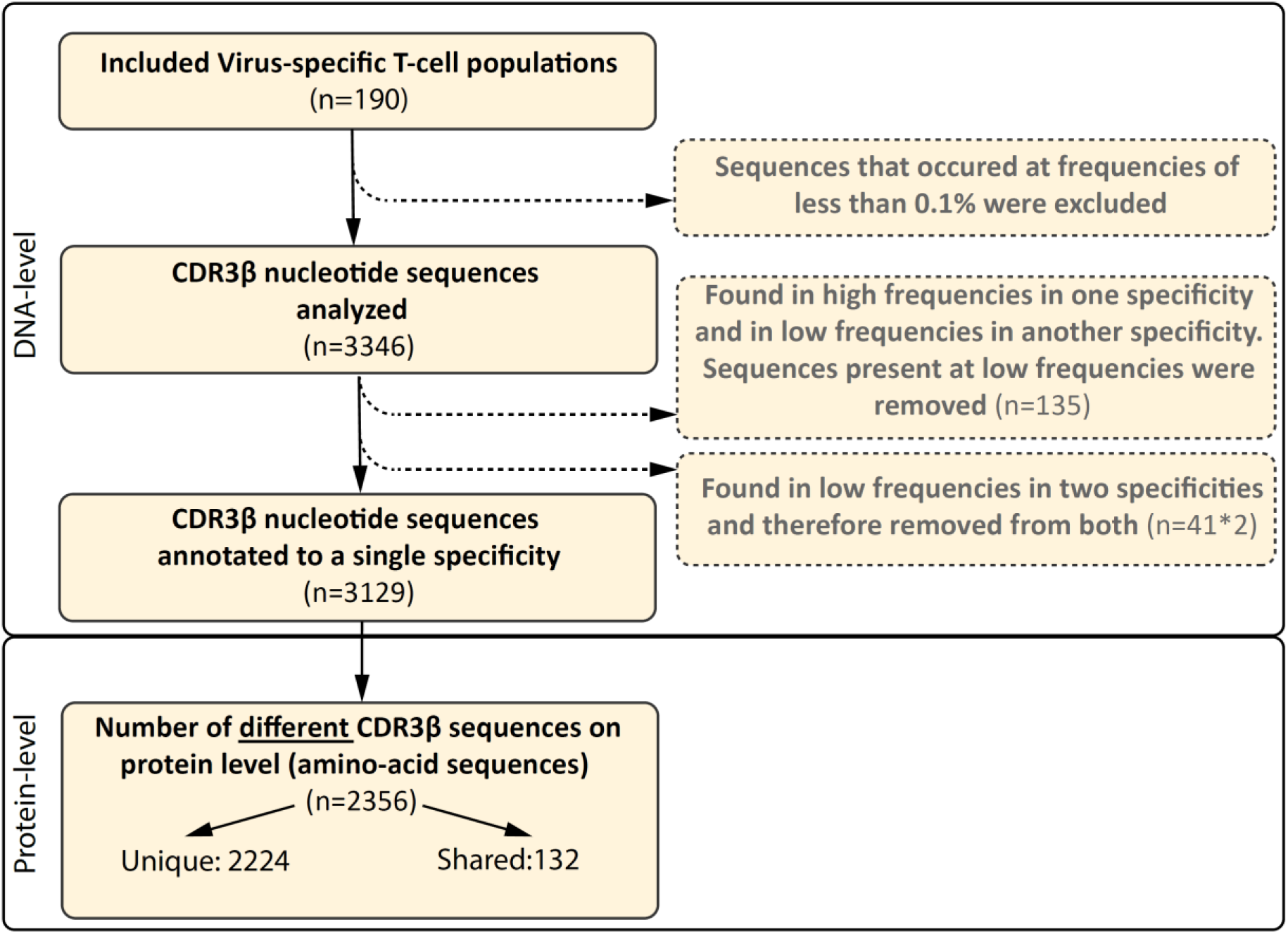
Flowchart of included and excluded CDR3β nucleotide and AA-sequences. In total, 190 different virus-specific T-cell populations were FACsorted using pMHC-tetramers, followed by a short-term *in vitro* stimulation. The CDR3β nucleotide-sequences were determined using next-gen Illumina sequencing. CDR3β nucleotide-sequences that occurred at a frequency of less than 0.1% in each sample were excluded. CDR3β nucleotide-sequences that were identical and present in two different specificities, but present at high frequencies in one specificity, were only removed from the specificity that contained the sequences at very low frequencies (0.1-0.5%; n=135). CDR3β nucleotide-sequences that were identical and present in two different specificities at low frequency were considered contamination and removed from the library (82 sequences, 41 different-sequences). The numbers of different CDR3β AA-sequences that were encoded by the CDR3β nucleotide-sequence are shown at protein level. We then assessed how many CDR3β-AA-sequences were found in multiple individuals (shared) and how many were only found in a single individual (unique).

To investigate the relationship between the numbers of CDR3β nucleotide-sequences and the translated number of CDR3β AA-sequences, we compared the nucleotide-sequences of the 131 PUB-I CDR3β AA-sequences found in different individuals. Different nucleotide-sequences can result in the same CDR3β AA-sequence, a phenomenon known as convergent recombination. We found that PUB-I CDR3β AA-sequences present at high (representative example**; Figure 2A**) or low frequencies (range 0.1-1%) (representative example; **Figure 2B**) could be encoded by different CDR3β nucleotide-sequences in the junctional regions of the CDR3β-regions in TCRs of T cells isolated from different individuals (**Figure 2C and 2D; Supplementary Figure 2**). Because the majority of nucleotide-sequences encoding the same CDR3β AA-sequences were different between individuals, these data exclude contamination as an explanation for the finding of PUB-I CDR3β AA-sequences.

**Figure 2.**
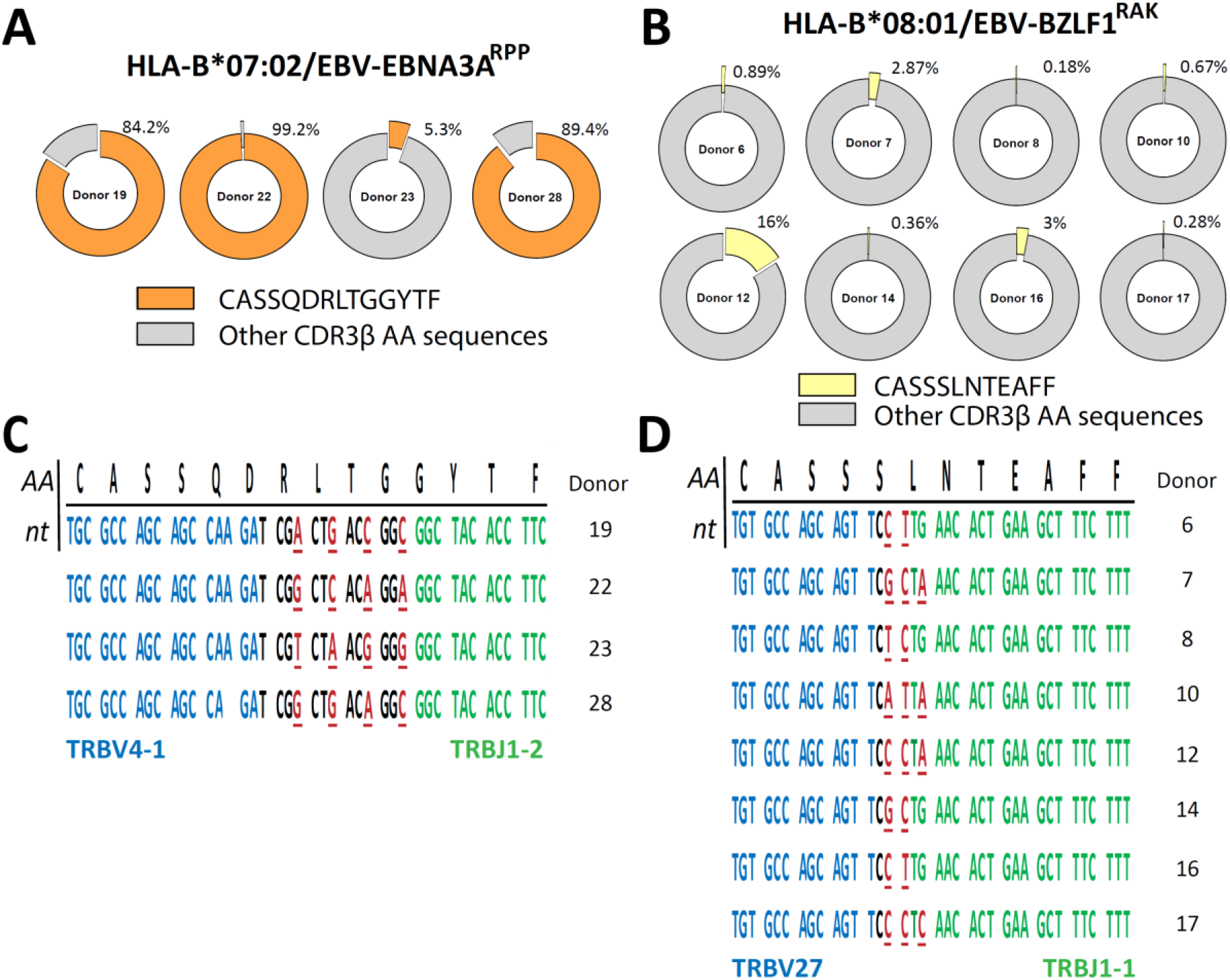
PUB-I CDR3β-sequences can be found in different individuals with small nucleotide-differences as a result of convergent recombination. The library of virus-specific CDR3β AA-sequences contained 131 sequences that were found in multiple different individuals (?2). The nucleotide-sequences of these CDR3β AA-sequences were analyzed to investigate differences and similarities. **A and B**) Shown are two representative examples of the frequencies of the PUB-I CDR3β AA-sequences CASQDRLTGGYTF and CASSSLNTEAFF, that were specific for HLA-B*07:02-restricted EBV-EBNA3A^RPP^ and HLA-B*08:01-restricted EBV-BZLF1^RAK^, respectively. **C and D**) Shown are the nucleotide-sequences of the CDR3β AA-sequences CASSQDRLTGGYTF and CASSSLNTEAFF that were shared by different individuals. Underlined nucleotides represent differences between the different individuals. Nucleotides in red represent differences to the consensus sequence. Nucleotide-sequences in blue and green represent perfect alignment with the germline sequences of the TRBV-gene and TRBJ-gene, respectively. AA; amino-acids, nt; nucleotides

### PUB-I and PUB-HS CMV-, EBV- and AdV-specific CDR3β AA-sequences are abundant in virus-specific T-cell populations

We then investigated the distribution of the 131 PUB-I CDR3β AA-sequences within the 21 different specificities and the prevalence among individuals for each of the PUB-I CDR3β AA-sequences per viral-antigen. T cells with PUB-I CDR3β AA-sequences were found for 19 out of the 21 specificities (**Supplementary Table 1**). PUB-I CDR3β AA-sequences were not observed in AdV-IE1^LLD^ and EBV-LMP2^ESE^-specific T-cell populations. Some T-cell populations (e.g. EBV-LMP2^FLY^) contained many different PUB-I CDR3β AA-sequences (n=24) that were all highly-similar. For this reason, we investigated the distribution of PUB-I CDR3β AA-sequences with unique TRBV and TRBJ-gene usage. This resulted in 29 different PUB-I CDR3β AA-sequences, distributed over 19 specificities **(Figure 3A; grey bars).** Six specificities contained two or three different (expressing different TRBV and/or TRBJ-genes) PUB-I CDR3β AA-sequences that were highly prevalent among individuals. To investigate how often these PUB-I CDR3β AA-sequences could be found in our cohort of healthy donors, we quantified the prevalence of each of these 29 PUB-I CDR3β AA-sequence **(Figure 3A; grey bars)**. Because we classified a PUB-I CDR3β AA-sequence as being present in at least 2 individuals, the prevalence among donors could not be less than 2 out of 17 (12%; 17 is maximum number of T-cell populations for 1 specificity). Only 4 out of 29 PUB-I CDR3β AA-sequences were found in only 2 individuals **(Figure 3A; grey bars)**. Overall, these 29 PUB-I CDR3β AA-sequences had a prevalence of 33% among healthy individuals (median; range 12%-82%). Importantly, most PUB-I CDR3β AA-sequences were found in at least 25% of individuals and 5 were even present in more than half the donors.

**Figure 3.**
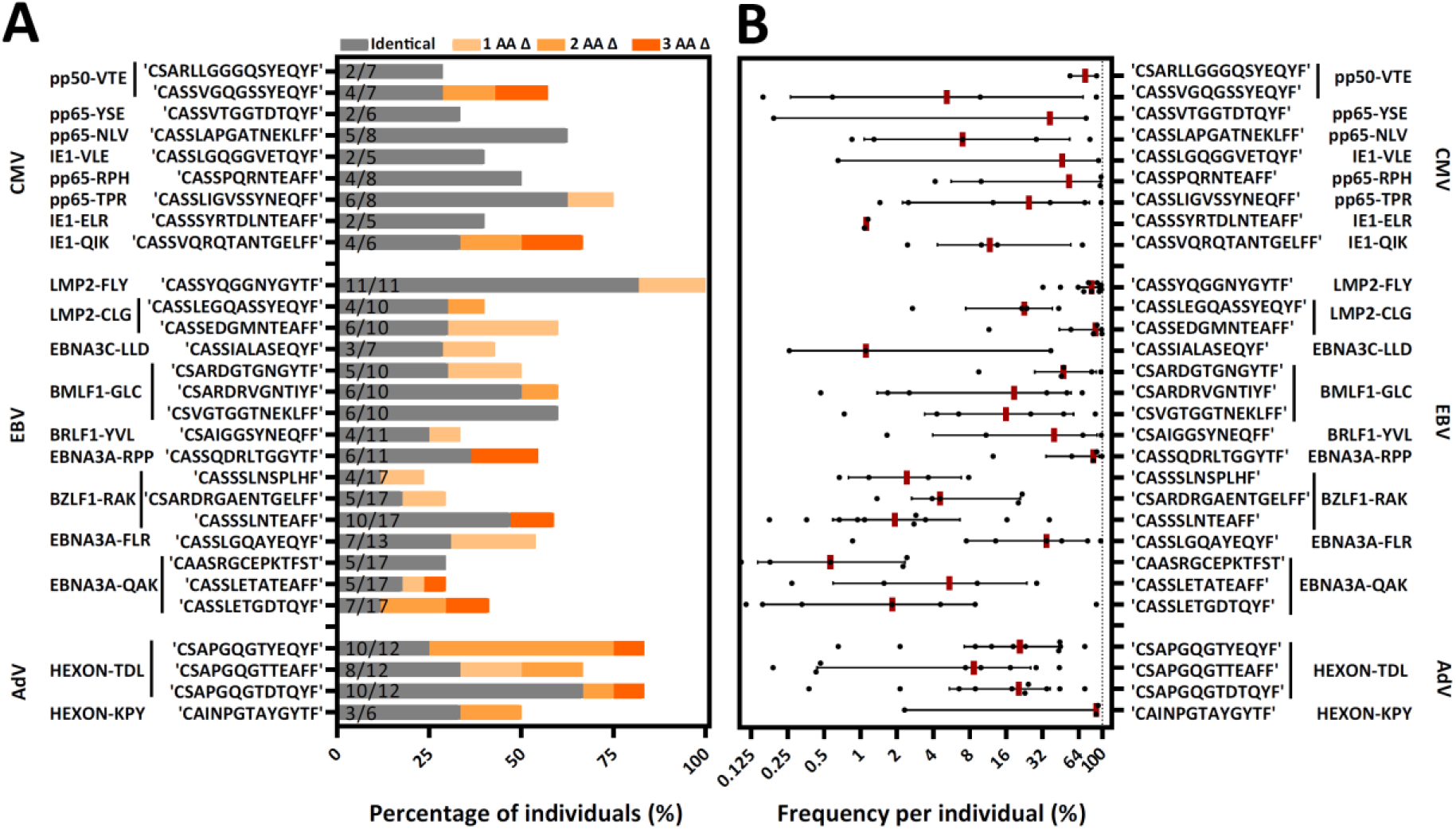
PUB-I and PUB-HS CMV, EBV and AdV-specific CDR3β AA-sequences are common in different individuals and of high frequencies within the T-cell specificities. **A)** Nine-teen different specificities contained PUB-I CDR3β-sequences, shared by at least 2 individuals. Six specificities contained 2 or 3 different (with different TRBV and/or TRBJ-genes) CDR3β AA-sequences that were found frequently in different individuals. The total numbers of different T-cell populations (different donors) that contained the PUB-I or PUB-HS CDR3β AA-sequences are indicated at the inner-side of the y-axis. The occurrence, shown as percentage among healthy individuals, is shown per CDR3β AA-sequence. PUB-I sequences are shown in grey. Individuals containing PUB-HS sequences with 1, 2 or 3 AA-differences (shown in light-dark orange) compared to the PUB-I sequence were stacked on top of the individuals where the PUB-I sequence was already identified. **B)** Shown is the sum of frequencies of the PUB-I and PUB-HS (1, 2 and 3 AA-differences) CDR3β AA-sequences per individual. Each dot is one individual, and the red-lines represent the medians with interquartile ranges. AA; amino-acids, nt; nucleotides, Δ; difference(s)

We and others hypothesized that the binding/docking of TCRs to HLA-peptide complexes might allow for small changes/flexibility in the CDR3 AA-sequences without significantly changing the conformation or interaction(23). Therefore, we investigated if there were CDR3β AA-sequences present in our data set that were highly-similar (PUB-HS) to PUB-I CDR3β AA-sequences and differed by 1, 2 or 3 AAs. The 2224 unique TCR nucleotide-sequences identified in our previous analysis may contain PUB-HS CDR3β AA-sequences that are in fact part of the same public response as the respective PUB-I TCRs. In total, 379 PUB-HS CDR3β nucleotide-sequences were present that also resulted in 379 PUB-HS CDR3β AA-sequences that differed by 1, 2 or 3 AAs from one of the 131 PUB-I CDR3β AA-sequences. This shows that 41% of the total virus-specific TCR-repertoire contained PUB-I and PUB-HS CDR3β nucleotide-sequences. We investigated if these PUB-HS CDR3β AA-sequences were also present in individuals that did not contain the respective PUB-I CDR3β AA-sequences. PUB-HS CDR3β AA-sequences were present for 21 out of 29 PUB-I CDR3β AA-sequences (**Figure 3A; shaded orange bars**). When we include the PUB-HS CDR3β AA-sequences and quantified the 29 PUB-I and PUB-HS CDR3β AA-sequences, these had a median prevalence of 50% among healthy individuals (range 23%-100%). The AdV-IE1^LLD^ and EBV-LMP2^ESE^-specific T-cell populations, where PUB-I CDR3β AA-sequences were not found, did contain PUB-HS CDR3β AA-sequences in multiple individuals at high frequencies(24) (**Supplementary Figure 3**). The frequencies of PUB-I combined with PUB-HS CDR3β AA-sequences were relatively high within each virus-specific T-cell population of each individual (**Figure 3B**). The frequencies of all PUB-I and PUB-HS CDR3β AA-sequences ranged from 0.1%-99.4% within the 19 different virus-specific T-cell populations with a median of 13.1%. When combined, all but one PUB-I plus PUB-HS CDR3β AA-sequences were found in at least 25% of individuals and 3 were even found in over 75% of individuals. These data show that for many PUB-I CDR3β AA-sequences we found sequences that were similar (1, 2 or 3 AA-differences), making up more than 40% of the total virus-specific TCR-repertoire and together these sequences were found in the majority of individuals at high frequencies.

### Identical and highly-similar CDR3β AA-sequences contain conserved regions in the junctional region

To investigate how the PUB-HS CDR3β AA-sequences related to the PUB-I CDR3β AA-sequences, we analyzed if the PUB-HS (1, 2 or 3 AA-differences) CDR3β AA-sequences showed variations at random positions or at specific positions compared to the PUB-I CDR3β AA-sequences. We hypothesized that if the binding/docking of PUB-HS TCRs was not significantly different, conserved regions and regions that allow for some variation could be identified in the CDR3β AA-sequences. As Illustrated in **Figure 4A** for the EBV-LMP2^FLY^-specific PUB-I CDR3β AA-sequence CASSYQGGNYGYTF, two motifs were identified with AA-differences predominantly located at positions 5 and 9/10 of the CDR3β-region. The AAs [QGG] at positions 6-8 were conserved for both motifs. In total, the two motifs consisted of 86 PUB-HS CDR3β AA-sequences with 1 or 2 AA-differences. The majority (n=71) had the same CDR3 length of 14 AAs as the PUB-I CDR3β AA-sequence, implying that variations were caused by AA-substitutions. The remaining 15 PUB-HS CDR3β AA-sequences had a CDR3 length of 15 AAs, due to AA-inserts, compared to the PUB-I CDR3β AA-sequence (**Figure 4B)**. Similar rules were found for the other 20 PUB-I CDR3β AA-sequences. Also here, some AA-positions were highly conserved, whereas others were variable. However, the precise locations of the variable AAs differed between specificities (Representative examples; **Figure 4C**).

**Figure 4.**
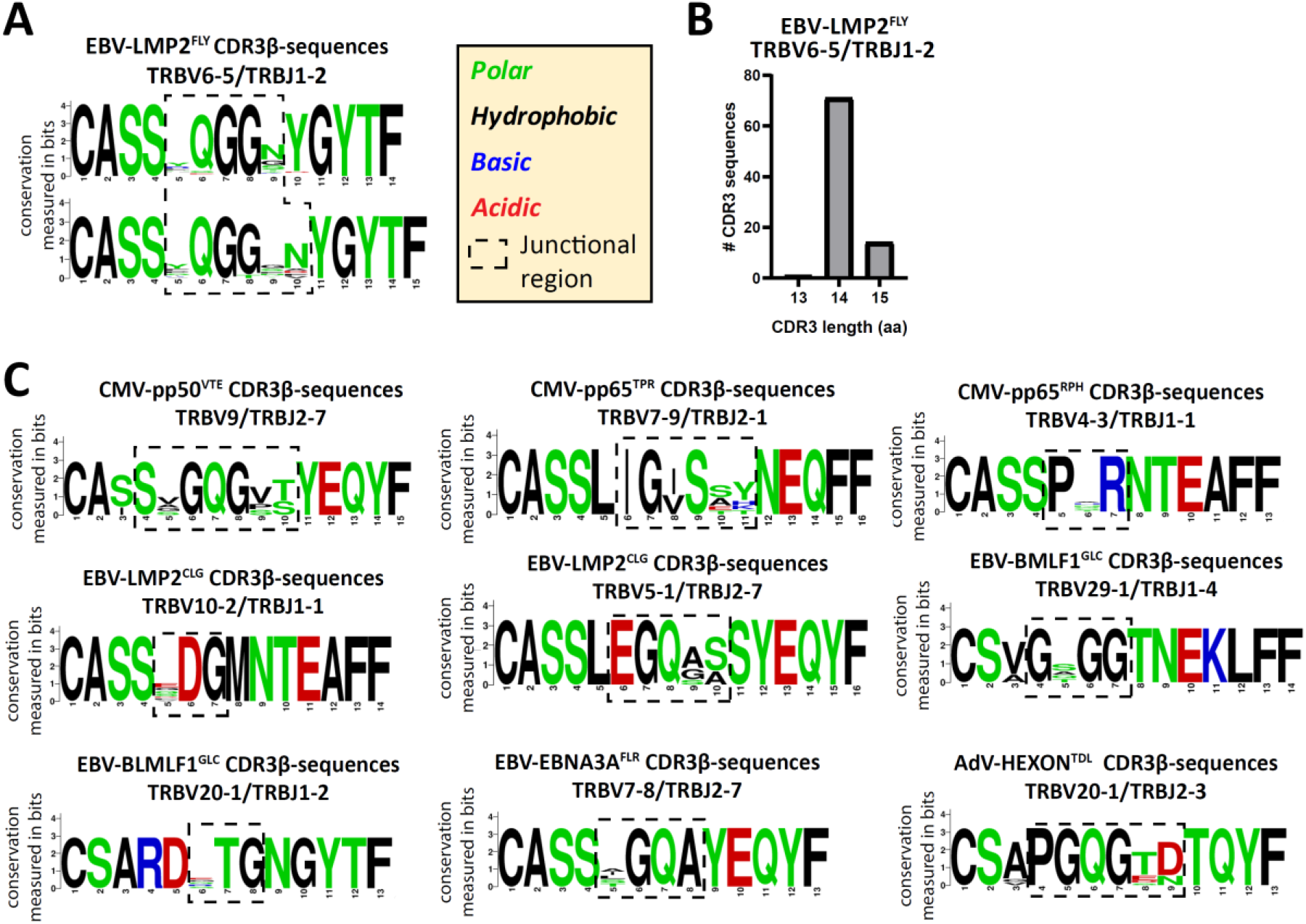
PUB-HS CDR3β AA-sequences show conserved regions and regions with high variability. The levenshtein distance was calculated (i.e. substitution, deletions or insertions of AAs) between CDR3β AA-sequences that express the same TRBV/TRBJ-genes as the PUB-I CDR3β AA-sequence. PUB-HS CDR3β AA-sequences were included with 1, 2 and 3 AA-differences compared to the PUB-I CDR3β AA-sequence. Sequence logos generated using WebLogo (http://weblogo.berkeley.edu/logo.cgi) show the relative frequency of each AA at each given position. The junctional regions (AAs that do not align with the germline TRBV or TRBJ-gene) are shown within the dotted-line box. **A)** The CDR3β AA-sequences specific for HLA-A*02:01-restricted EBV-LMP2^FLY^ with a CDR3 length of 14 and 15 AAs were stacked and show conserved and variable regions in the CDR3β-region. **B)** Shown are the CDR3 length distributions of CDR3β AA-sequences specific for EBV-LMP2^FLY^-expressing TRBV6-5/TRBJ1-2. **C)** Shown are representative examples of PUB-HS (1, 2 or 3 AA differences) and PUB-I CDR3β AA-sequences that were stacked per specificity and TRBV/TRBJ-usage.

As a control, we assessed if these conserved motifs were predictive for the specificity when searching in our database of 2355 unique CDR3β AA-sequences. The requirement was that each motif should not be present in another specificity. We observed that some specificities contained motifs of only 3 or 4 AAs that were exclusive for that specificity and were not observed in any other specificity **(Table 3).** Altogether, these data show that the variations in the CDR3β AA-sequences were not random, but occurred at specific positions that resulted in conserved regions that were predictive for the specificities.

**Table 3.**
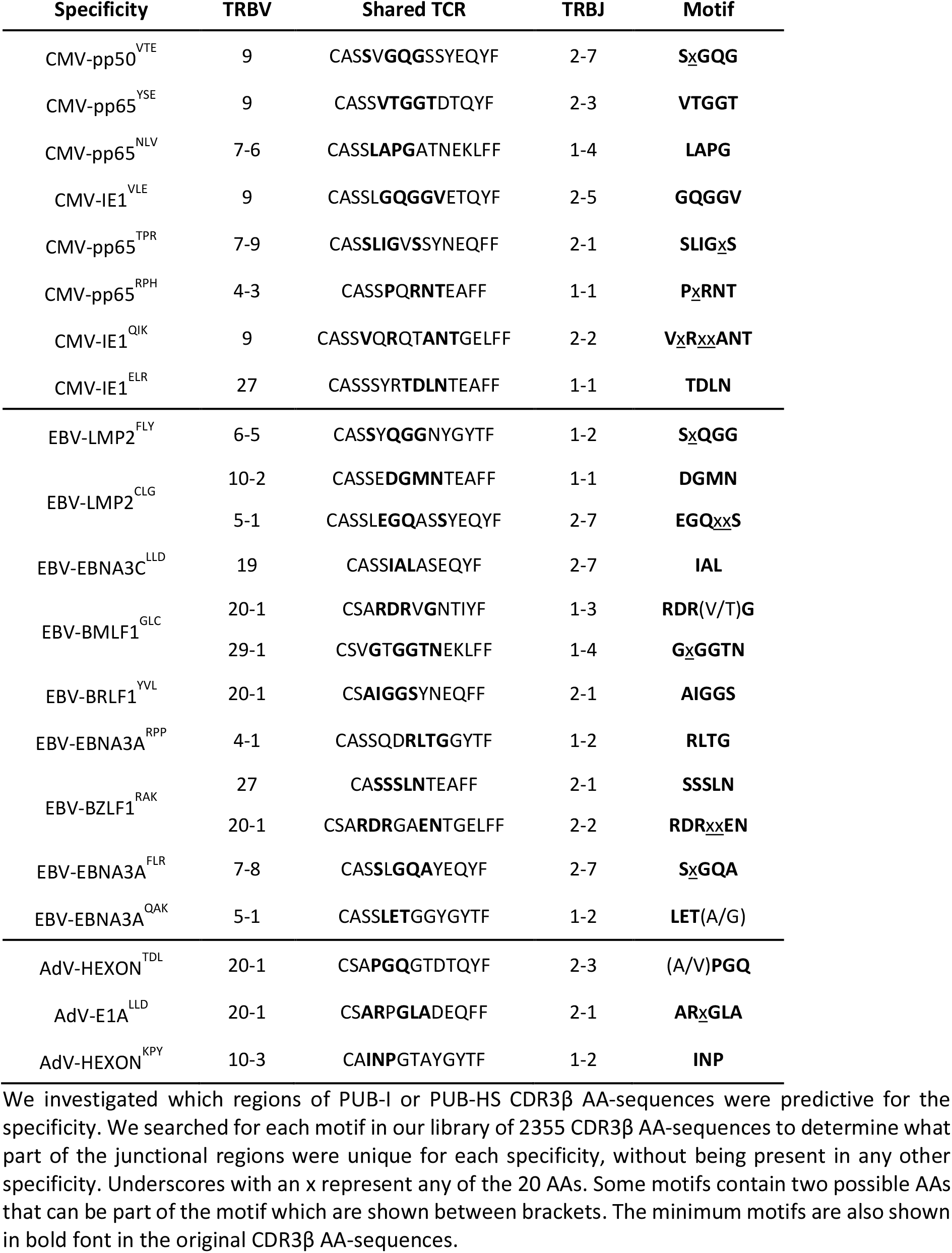
Conserved motifs that predict the specificity.

### Computational analysis reveals conserved regions in CDR3β AA-sequences despite using different TRBJ-genes

We hypothesized that if the conserved junctional region is a crucial part of the peptide-HLA binding, virus-specific TCR-repertoires could also contain CDR3β AA-sequences with the same conserved region, while allowing different TRBJ-gene usage, as long as the 3-dimensional conformation would allow this. Since the TRBJ-regions often differ by more than 3 AAs, we were not able to include these as PUB-HS CDR3β AA-sequences. Such PUB-HS CDR3β AA-sequences that use different TRBJ-genes might even further increase the prevalence of PUB-I and PUB-HS CDR3β AA-sequences in the virus-specific TCR-repertoire. To investigate this, we performed a computational analysis using the levenshtein-distances (AA-differences) between all different CDR3β AA-sequences. For four different specificities (EBV-LMP2^FLY^, EBV-EBNA3A^RPP^, AdV-E1A^LLD^, and AdV-HEXON^TDL^) we observed clustering of CDR3β AA-sequences that expressed the same TRBV-genes while using different TRBJ-genes. For example (**Figure 5A**), the HLA-A*02:01-restricted EBV-LMP2^FLY^-specific CD8^pos^ T-cell repertoire contained 2 clusters within the cluster of TRBV6-5-expressing T cells (TRBV6-5/TRBJ1-2 and TRBV6-5/TRBJ2-1). The majority of CDR3β AA-sequences within the TRBJ1-2 cluster had a length of 14 AAs, while CDR3β AA-sequences from the TRBJ2-1 cluster had a length of 13 AAs **(Figure 5B)**. Analysis of the junctional regions of the TRBV6-5/TRBJ1-2 and TRBV6-5/TRBJ2-1-encoded CDR3β AA-sequences revealed strong conservation of AAs [QGG] on positions 6-8, despite different TRBJ-usage and CDR3 lengths **(Figure 5C)**. Similarly, the HLA-A*01:01-restricted AdV-HEXON^TDL^-specific CD8^pos^ T-cell repertoire contained two large clusters of CDR3β AA-sequences, using TRBV20-1 or TRBV5-1 (**Figure 5D**), all with a CDR3 length of 13 AAs. The first cluster (TRBV20-1) contained sub-clusters of CDR3β AA-sequences using TRBJ1-1, TRBJ2-3 or TRBJ2-7 and the second cluster (TRBV5-1) contained CDR3β AA-sequences using TRBJ2-1 or TRBJ2-7. AdV-HEXON^TDL^-specific CDR3β AA-sequences expressing TRBV20-1 revealed strong conservation of AAs [PGQG] on positions 4-7, which fell outside the region encoded by TRBJ (**Figure 5E**). Additionally, AdV-HEXON^TDL^-specific CDR3β AA-sequences expressing TRBV5-1 revealed strong conservation of AAs [N__D] on positions 4 and 7, despite different TRBJ-usage. These examples illustrate that virus-specific TCR-repertoires can have conserved CDR3β-regions, while using different TRBJ-genes, allowing substantial variability at specific positions encoded by the TRBJ-region. This will further increase the prevalence of PUB-I and PUB-HS CDR3β AA-sequences in the total virus-specific TCR-repertoire.

**Figure 5.**
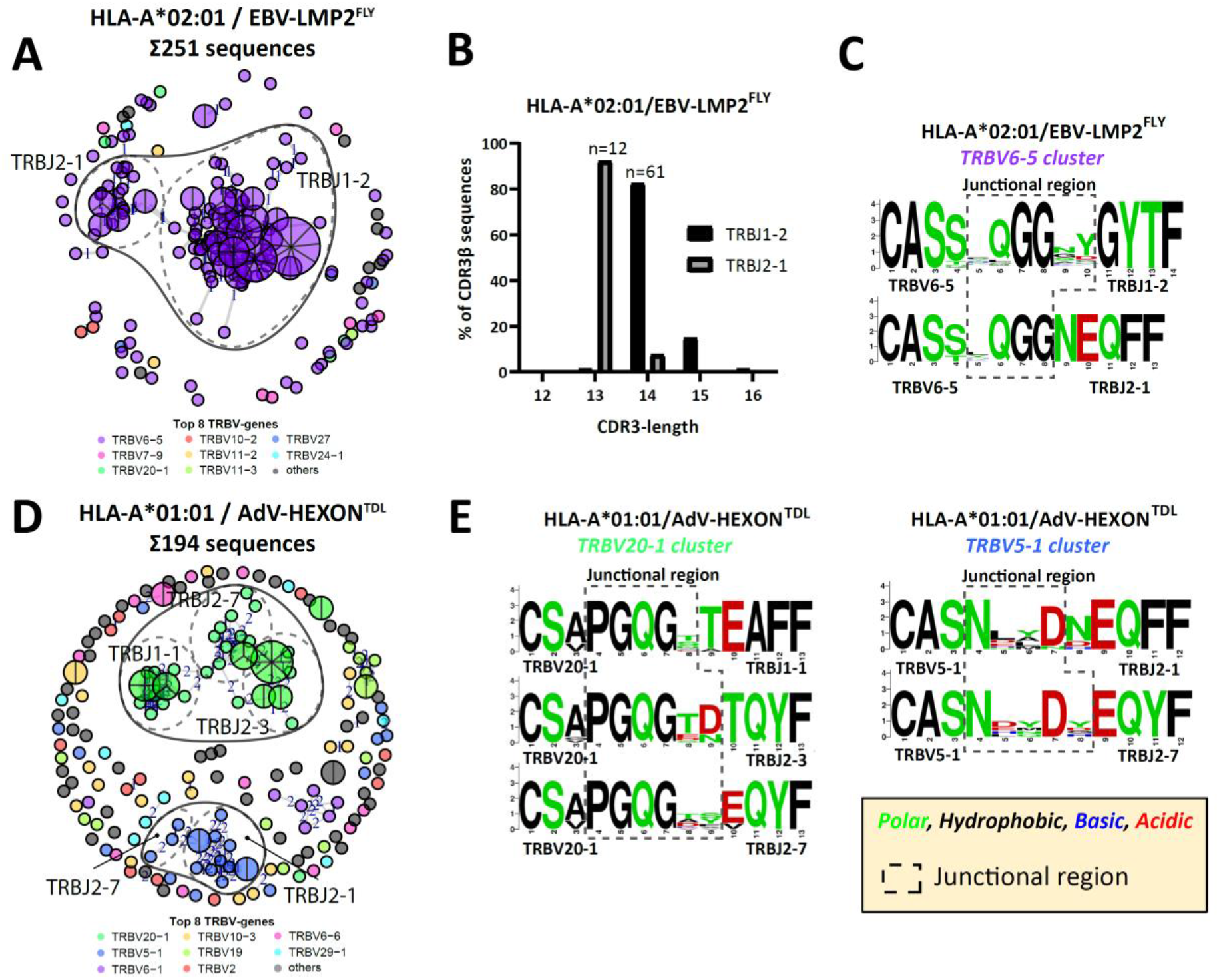
Computation analysis reveals clustering of PUB-HS CDR3β AA-sequences that contained conserved regions in the CDR3β-region. Computational analysis was performed using the levenshtein distance (differences in AAs) between CDR3β AA-sequences of one specificity. CDR3β AA-sequences were plotted and colored according to the top 8 most frequent TRBV-genes and were clustered and linked by a line if they were similar, with a number (levenshtein distance of 1, 2 or 3) representing the differences in AAs. PUB-I CDR3β AA-sequences were plotted as a pie-chart, whereby the size and number of slices indicate in how many individuals this CDR3β AA-sequence was present. Sequence logos generated using WebLogo (http://weblogo.berkeley.edu/logo.cgi) show the relative frequency of each AA at each given position. The junctional region (AAs that do not align with the germline TRBV or TRBJ-gene) are shown within the box with the dotted-line **A**) Shown is a representative example of a virus-specific CD8^pos^ T-cell population, specific for EBV-LMP2^FLY^ with overlapping clusters of sequences that express different TRBJ-genes, while expressing the same TRBV-gene. **B**) The lengths of the CDR3β-regions of the two clusters of EBV-LMP2^FLY^-specific CDR3β-sequences are shown. Varying lengths of the CDR3β-region within a cluster would suggest deletions or insertions, whereby the same length would indicate AA substitutions. **C**) Shown are the sequence motifs of the two EBV-LMP2^FLY^-specific clusters. **D)** Shown is a second representative example of a virus-specific CD8^pos^ T-cell population, specific for AdV-HEXON^TDL^, with overlapping clusters of sequences that express different TRBJ-genes, while expressing the same TRBV-gene. **E)** Shown are the sequence motifs of the TRBV20-1 and TRBV5-1-expressing EBV-HEXON^TDL^-specific clusters.

### Individuals with heterozygous HLA backgrounds contain the same shared identical and highly-similar CDR3β AA-sequences

To determine whether the magnitude of PUB-I and PUB-HS CDR3β AA-sequences was particular for our cohort of individuals with a homozygous HLA background, we investigated if the same phenomenon was also present in individuals with a heterogeneous HLA background. We performed the same analyses on virus-specific CD8^pos^ T-cell populations targeting 11 different viral antigens that were generated and used in the context of a clinical study(25). A total of 1157 CDR3β nucleotide-sequences could be correctly annotated. In total, 695 (61%) nucleotide-sequences resulted in unique CDR3β AA-sequences, that were only found in one individual, and 462 nucleotide-sequences (39%) resulted in 89 different PUB-I CDR3β AA-sequences. From the 695 unique CDR3β AA-sequences, 134 PUB-HS CDR3β nucleotide-sequences were present that differed by 1, 2 or 3 AAs from one of the 89 PUB-I CDR3β AA-sequences. This shows again that also in this cohort a large part (51%) of the total virus-specific TCR-repertoire contained PUB-I and PUB-HS CDR3β nucleotide-sequences. Because the targeted viral antigens were not fully identical in both cohorts, we could investigate the prevalence of 20 out of 29 PUB-I CDR3β AA-sequences in this cohort. In total, 17 out of 20 CDR3β AA-sequences that were previously identified, could also be identified in this independent cohort. When we included the PUB-HS CDR3β AA-sequences and quantified the 17 PUB-I and PUB-HS CDR3β AA-sequences, these sequences had a similar high prevalence of a median of 89% among healthy individuals (range 26-100%). (**Figure 6A).** These CDR3β AA-sequences were also present at high frequencies within each virus-specific T-cell population (**Figure 6B**). These data show that the same PUB-I or PUB-HS CDR3β AA-sequences are also present in virus-specific T cells isolated from an independent cohort of individuals with a heterogeneous HLA background with a similar prevalence among donors and frequency within donors.

**Figure 6.**
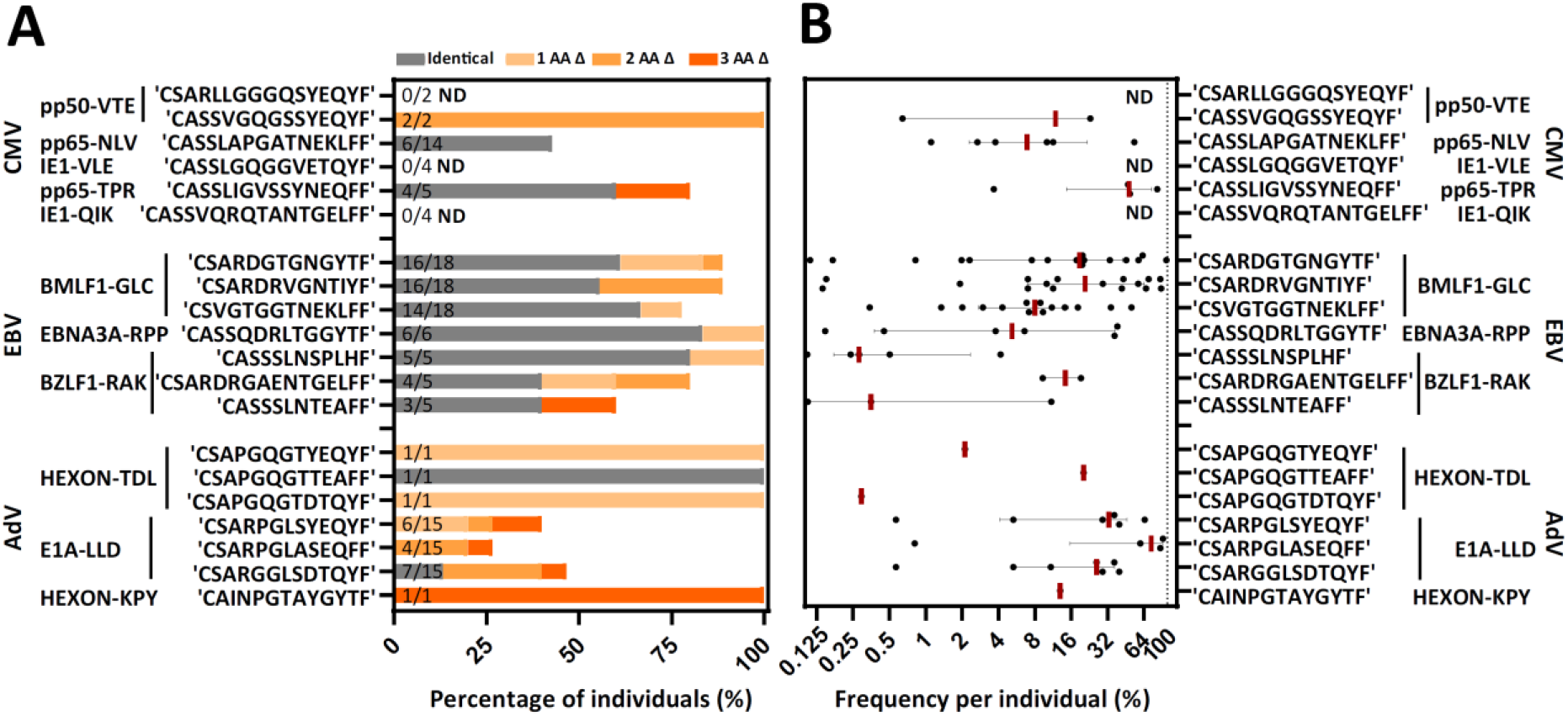
Individuals with a different HLA background from an independent database contain the same PUB-I and PUB-HS CDR3β AA-sequences. Virus-specific T-cell populations targeting 11 different viral-antigens, derived from an independent database, could be evaluated for the occurrence of PUB-I pr PUB-HS CDR3β AA-sequences. **A)** Nine out of the 11 specificities contained the same PUB-I or PUB-HS CDR3β AA-sequences as in our database. The occurrence, shown as percentages among healthy donors, is shown per CDR3β AA-sequence. PUB-I sequences are shown in grey. PUB-HS CDR3β AA-sequences from other individuals with 1, 2 or 3 AA differences, were stacked on top of the already identified public-identical sequences. The total numbers of different T-cell populations (different donors) that contained the PUB-I or PUB-HS CDR3β-sequences are indicated at the inner-side of the y-axis. **B)** Shown are the sum of frequencies of the PUB-I or PUB-HS CDR3β AA-sequences per donor. Each dot is one donor shown and the red-lines represent the medians with interquartile ranges. AA; amino-acid, Δ; difference(s), ND; Not detected

## Discussion

In this study, we quantitatively analyzed the magnitude, defined as prevalence within the population and frequencies within individuals, of public-identical (PUB-I) together with public-highly-similar (PUB-HS) TCRs in TCR-repertoires of CMV, EBV and AdV-specific CD8^pos^ T-cell populations. In total, 2224 (71%) TCR-CDR3β nucleotide-sequences resulted in unique CDR3β AA-sequences, and 905 nucleotide-sequences (29%) resulted in 131 different PUB-I CDR3β AA-sequences that were found in two or more unrelated individuals. These PUB-I CDR3β AA-sequences were distributed over 19 out of 21 virus-specificities and contained 29 different PUB-I CDR3β AA-sequences that were often found in multiple individuals at high frequencies. The virus-specific T-cell populations additionally contained 12% PUB-HS CDR3β AA-sequences, which differed by 1, 2 or 3 AAs compared to the respective PUB-I CDR3β AA-sequences. PUB-HS CDR3β AA-sequences could be found in virus-specific T-cell populations of individuals who did not contain the PUB-I CDR3β AA-sequence as well as of individuals who already contained the PUB-I CDR3β AA-sequence. Analysis of the PUB-I and PUB-HS CDR3β AA-sequences revealed strong conservation of specific AA motifs in the junctional region together with variability of AAs at specific positions at the TRBV/TRBD- and/or TRBD/TRBJ-border regions. Positions with high variability were often adjacent to or even interspersed with the conserved motif. The conserved motifs that we identified were unique for each specificity, and could not be identified in any other specificity in our database. This makes it very likely that these motifs are important for binding of the TCRs to the peptide-HLA complexes. Combined, 41% of the total virus-specific TCR-repertoire consisted of PUB-I and PUB-HS CDR3β nucleotide-sequences. These findings were based on virus-specific T-cell populations derived from two homogeneous donor cohorts that homozygously expressed HLA-A*01:01/HLA-B*08:01 or HLA-A*02:01/HLA-B*07:02. However, we found similar high percentages (51%) of PUB-I and PUB-HS CDR3β nucleotide-sequences within virus-specific T-cell populations from healthy donors with heterogeneous HLA-backgrounds that were generated for a recent clinical study(25). These dominant PUB-I and PUB-HS TCRs probably are a reflection of the viral-antigen-specific T-cell responses that most optimally encountered the peptide-HLA complexes on the infected target cells and could be utilized for the design of future immunotherapy purposes including TCR-gene transfer strategies.

Various explanations have been suggested to underlie the development of public TCRs in T-cell responses targeting the same antigenic epitope(26). One was a high probability that these PUB-I sequences can be generated during V-D-J recombination(27, 28). Furthermore, various nucleotide-sequences can result in the same TCR AA-sequences that further increase the probability(29). Selection *in vivo* by optimal antigen-specific proliferation may result in a dominant antigen-specific memory T-cell population(30). These determinants may also lead to TCRs that are highly-similar to the PUB-I sequence, although they were often not included in the analyses of such public T-cell responses. It has been shown that conserved AAs in the CDR3 loop provide a structural framework that is required for the maintenance of the three dimensional TCR-structure(31). A similar structural framework between the PUB-I and PUB-HS sequences can thus lead to a conserved engagement with the peptide/HLA complex(32). Our rationale is that the PUB-I and PUB-HS sequences are part of the same public T-cell response when the same peptide-HLA complex is targeted, the same variable gene is expressed to have identical CDR1 and CDR2 regions and contains the same conserved AAs in the CDR3 loop. With this set of rules, we were able to quantitatively analyze the public T-cell responses and showed that T cells expressing PUB-I TCRs together with T cells that express PUB-HS TCRs made up at least 41% of the total TCR-repertoire.

These percentages of PUB-I and PUB-HS TCRs within these virus specific T-cell responses may still be an underestimation since the prerequisite of the identification of a PUB-HS TCR was similarity to a PUB-I TCR that was present in at least 2 individuals. Highly similar TCRs with only mutual similarities without identity in at least 2 individuals were not included as PUB-HS TCRs. Therefore, some of the unique TCRs within the virus-specific T-cell repertoire may also be part of a public T-cell response. This was indeed illustrated by the growing percentages of PUB-I and PUB-HS sequences when including more sampled sequences(33). Although it was suggested that HLA polymorphisms might be a confounding factor that affect the sharing of TCRs(33, 34), we showed that our validation cohort with different HLA-backgrounds revealed frequencies of the PUB-I and PUB-HS TCRs with at least a similar magnitude.

In conclusion, our findings demonstrate that a large part of the virus-specific TCR-repertoire contains PUB-I and PUB-HS TCRs at high frequencies in multiple different individuals. Because T cells that express these TCRs apparently proliferated and differentiated into memory T cells, it is highly likely that they participated in control of the virus. Since it is plausible that the highly-similar TCRs with conserved motifs similarly dock to the peptide-HLA complex as the identical shared TCR-sequences, these PUB-I and PUB-HS sequences can be considered part of the same public T-cell response. Such public TCRs may be used for therapeutic benefit in TCR-gene transfer-based immunotherapy strategies to effectively control viral-reactivation in immunocompromised patients.

## Methods

### Collection of donor material

After informed consent according to the Declaration of Helsinki, healthy individuals (homozygously) expressing HLA-A*01:01 and HLA-B*08:01 or HLA-A*02:01 and HLA-B*07:02 were selected from the Sanquin database and the biobank of the department of Hematology, Leiden University Medical Center (LUMC). Peripheral blood mononuclear cells (PBMCs) were isolated by standard Ficoll-Isopaque separation and used directly or thawed after cryopreservation in the vapor phase of liquid nitrogen. Donor characteristics (HLA typing, CMV and EBV serostatus) are provided in **Table 2.** For isolation of donor-derived virus-specific T cells using fluorescence-activated cell sorting (FACS, gating strategy see **supplementary figure 4**) with pMHC-tetramers (**Table 1**) and expansion of donor-derived virus-specific T cells, see supplementary material and methods.

### TCRβ-library preparation

TCRβ-sequences were identified using ARTISAN PCR adapted for TCR PCR(35, 36). Total mRNA was extracted from 190 pMHC-tetramer^pos^ purified (**Supplementary Figure 1A**) virus-specific T-cell populations(37) using magnetic beads (Dynabead mRNA DIRECT kit; Invitrogen, Thermo Fisher Scientific). Ten μl (~1μg) of mRNA per sample was mixed with TCRβ constant region-specific primers (1μM final concentration) and SmartSeq2modified template-switching oligonucleotide (SS2m_TSO; 1μM final concentration) and denatured for 3 minutes at 72°C. After cooling, cDNA was synthesized for 90 minutes at 42°C with 170 U SMARTscribe reverse transcriptase (Takara, Clontech) in a total volume of 20μl containing 1.7U/μl RNasin (Promega), 1.7mM DTT (Invitrogen, Thermo Fisher Scientific), 0.8mM each of high-purity RNAse-free dNTPs (Invitrogen, Thermo Fisher Scientific) and 4μl of 5x first-strand buffer. During cDNA synthesis, a non-templated 3’polycytosine terminus was added (**Supplementary Figure 1B**), which created a template for extension of the cDNA with the TSO(38). PCR (2min at 98°C followed by 40 cycles of [1s at 98°C, 15s at 67°C, 15s at 72°C], 2 min at 72°C) of 5μl of cDNA was then performed using Phusion Flash (Thermo Fisher Scientific) with anchor-specific primer (SS2m_For; 1μM final concentration) and each (1μM final concentration) of the nested primers specific for the constant regions of TCRβ constant 1 and TCRβ constant 2. Both forward and reverse PCR primers contained overhanging sequences suitable for barcoding. Amplicons were purified and underwent a second PCR (2min at 98°C followed by 10 cycles of [1s at 98°C, 15s at 65°C, 30s at 72°C], 2 min at 72°C) using forward and reverse primers (1μM final concentration) with overhanging sequences containing identifiers (sequences of 6 base-pairs) and adapter sequences appropriate for Illumina HiSeq platforms (or PacBio; Pacific Biosciences). Unique identifiers were used for each T-cell population targeting one antigen. Forward or reverse identifiers were shared between T-cell populations targeting different antigens. For all primer sequences see **Supplementary Table 2**. For identifier sequences see **Supplementary Table 3.** Amplicons with identifiers were purified, quantified and pooled into one library for paired-end sequencing of 150bp on an Illumina HiSeq4000. Deep sequencing was performed at GenomeScan (Leiden, The Netherlands). Raw data were de-multiplexed and aligned to the matching TRBV, TRBD, TRBJ and constant (TRBC) genes. CDR3β-sequences were built using MIXCR software using a bi-directional approach (5’-3’ and 3’-5’ read)(39). CDR3β-sequences with a stop-codon were removed from the library. Bi-directional readings using MIXCR could result in out-of-frame CDR3β AA-sequences due to the even number of nucleotides. These sequences (n=392) were manually aligned with the germline TRBV and TRBJ-sequence. CDR3β-sequences were further processed using custom scripts in R to compare specificities and sharing of CDR3β-sequences.

### Computational unbiased repertoire analysis

The following R-packages were used in R-software to generate a nodal plot of CDR3β AA-sequences with the levenshtein distance as parameter for similarity: “igraph” to create network objects, obtain the degree of a node and its betweenness(40), “data.table” to organize CDR3β-sequences; “stringdist” to calculate Levenshtein distances(41), “Biostrings” for fast manipulation of large biological sequences or sets of sequences(42), “dplyr” to arrange and filter data(43), “tibble” for providing opinionated data frames, “ggplot2” for generating figures(44) and “RColorBrewer” to create graphics(45). A levenshtein distance of 0.25 was added to visualize multiple identical sequences. Nodes with identical sequences (levenshtein distance of 0.25) were manually replaced by pie-charts using Adobe Illustrator CC 2018.

### Sequence logo plots

To identify which positions of PUB-I and PUB-HS CDR3β AA-sequences were conserved and which were variable, all CDR3β AA-sequences with the most frequent CDR3 length were included and the AAs were stacked for each position in the sequence. The overall height of the stacks indicates the sequence conservation at that position, while the height of symbols within the stacks indicates the relative frequency of each AA at that position. AAs have colors according to their chemical properties; polar AAs (G, S, T, Y, C, Q, N) show as green, basic (K, R, H) blue, acidic (D, E) red, and hydrophobic (A, V, L, I, P, W, F, M) AAs as black(46).

### Generation of independent TCR-database from virus-specific T-cell products generated for a clinical trial

As an independent TCR-database containing TCR-sequences from virus-specific T cells, we used the information obtained from the virus-specific T-cell products generated in the context of the phase I/II safety and feasibility study T Control (EudraCT-number 2014-003171-39) using the MHC-I-Streptamer isolation technology (Juno Therapeutics, Munich, Germany)(25, 47). Sequencing was performed as described above for all virus-specific T-cell populations per donor, resulting in unique identifiers for all virus-specific T-cell populations in the TCRβ-library.

### Data deposition

TCR-Sequencing data is deposited to the VDJdb (https://vdjdb.cdr3.net/) and will be available upon acceptance

## Acknowledgement

This work was supported by Sanquin Research and Landsteiner Laboratory for Blood Cell research [PPO 15-37/Lnumber 2101]. This study was in part also supported by research funding from Stichting den Brinker (The Netherlands, Zeist) that made a donation to the Leukemia fund from the Bontius Foundation (Leiden University Medical Center).

## Authorship

Contribution: W.H., D.A.T., L.H. and M.C.J.R performed experiments, K.A and G.A.E designed the R-script for computational analyses, W.H. analyzed results and made the figures, W.H., J.H.F.F., D.A. and I.J. designed the research and wrote the paper.

## Conflict-of-interest disclosure

The authors declare no competing financial interests.

## Correspondence

Wesley Huisman, Department of Hematology, Leiden University Medical Center, The Netherlands;

## Notes

**Potential conflict of interest**, The authors do not have a commercial or other association that might pose a conflict of interest.

### Competing Interest Statement

The authors have declared no competing interest.

