## Supplementary appendix for "Public T-Cell Receptors (TCRs) Revisited by Analysis of the Magnitude of Identical and Highly-Similar TCRs in Virus-Specific T-Cell Repertoires of Healthy Individuals"

**Online Supplementary Appendix**

**Material and Methods**

**Generation of peptide-MHC complexes to isolate virus-specific T cells**

All viral peptides were synthesized in-house using standard Fmoc chemistry. Recombinant HLA-A*01:01, HLA-A*02:01, HLA-B*07:02 and HLA-B*08:01 heavy chain and human β2m light chain were in-house produced in Escherichia coli. MHC-class-I refolding was performed as previously described with minor modifications[1]. Major histocompatibility complex (MHC)-class-I molecules were purified by gel-filtration using HPLC. Peptide-MHC(pMHC) tetramers were generated by labeling biotinylated pMHC-monomers with streptavidin-coupled phycoerythrin (PE; Invitrogen, Carlsbad, USA), allophycocyanin (APC, Invitrogen), brilliant violet 421 (BV421, Becton Dickinson (BD), Franklin Lakes, USA), brilliant violet 510 (BV510, BD) or peridinin-chlorophyll-protein complex (PerCP, Invitrogen). Complexes were stored at 4 °C. Formation of stable pMHC-monomers was performed using UVexchange technology[2] and according to a previously described protocol[3].

### Isolation and expansion of virus-specific T cells

Phycoerythrin (PE), allophycocyanin (APC), BV421, BV510 and/or peridinin-chlorophyll-protein (PerCP)-labeled pMHC-tetramer complexes were used for fluorescence-activated cell sorting (FACSorting). The pMHC-tetramers used are shown in **Table 2**. Per specificity, 30*$10^6$ PBMCs were first incubated with pMHC-tetramers at 4°C for 30 min, followed by labeling with APC-H7 CD8 (BD) and fluorescein isothiocyanate-labeled (FITC) CD4 and CD14 (BD) antibodies at 4°C for 30 min. PeptideMHC-tetramer positive, CD8$^{pos}$/CD4$^{neg}$ T cells were FACsorted and seeded at 10,000 cells per well in U-bottom microtiter plates for the generation of bulk T-cell populations. Peptide-MHC-tetramer$^{pos}$ virus-specific T cells, targeting a single antigen, were first specifically expanded in the presence of $10^{-7}$M of the specific peptide in T-cell medium: Iscove's Modified Dulbecco's Medium (IMDM; Lonza, Verviers, Belgium) containing 5% heat-inactivated fetal bovine serum (FBS; Invitrogen), 5% heat-inactivated human serum (ABOS; Sanquin Reagents, Amsterdam, The Netherlands), 100U/mL penicillin (Lonza), 100µg/mL streptavidin (Lonza) , 2.7mM L-glutamine (Lonza), and 100IU IL-2/mL (Chiron, Emeryville, USA) and with 5-fold 35 Gy irradiated autologous PBMCs as feeder cells. Initial specific stimulation and expansion with $10^{-6}$M peptide was performed to stimulate preferential outgrowth of pMHC-tetramer$^{pos}$ T cells. After two weeks of culture, pMHC-tetramer$^{pos}$ T-cell populations were qualified as pure populations when they contained ≥97% pMHC-tetramer$^{pos}$ cells. Sorting was performed on a FACS ARIA (BD) and analyzed using Diva software (BD). All analyses were performed on a FACS Calibur (BD), and analyzed using Flowjo Software (TreeStar, Ashland, USA).

**Results**

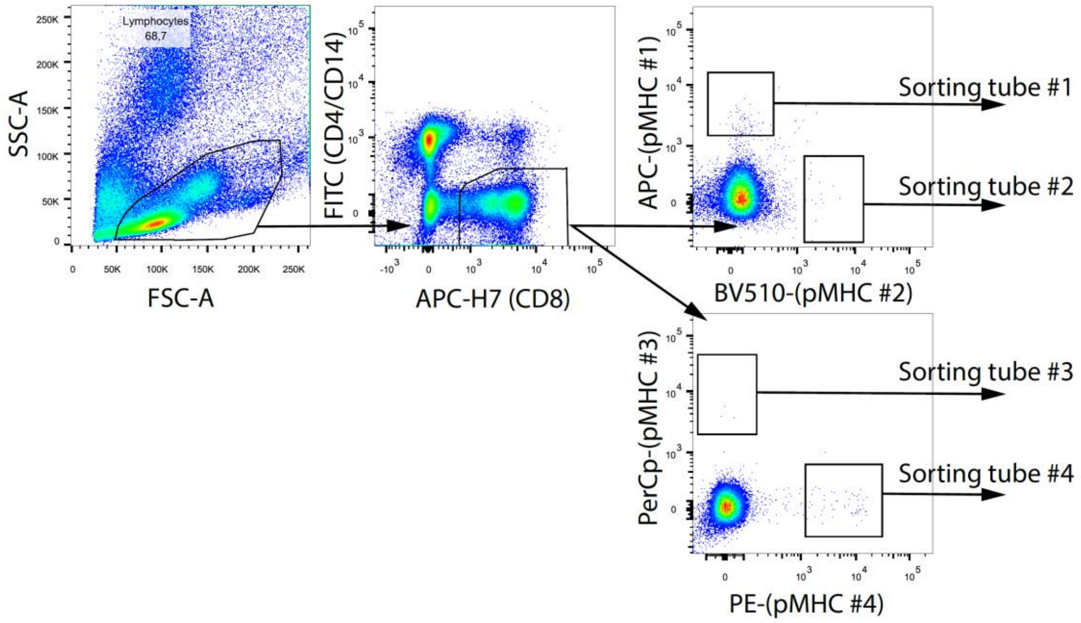

**Supplementary Figure 1: Gating strategy for a 4-way single-pMHC-tetramer sort from PBMCs.** In total, $30*10^6$ PBMCs were incubated with 4 different pMHC-tetramer complexes, followed by labeling with CD8, CD4 and CD14 monoclonal antibodies. Viable cells were gated based on FSC/SSC followed by gating of $CD8^{pos}$ and $CD4/CD14^{neg}$ T cells. Peptide-MHC-tetramer positive T cells were sorted simultaneously for 4 different specificities in bulk.

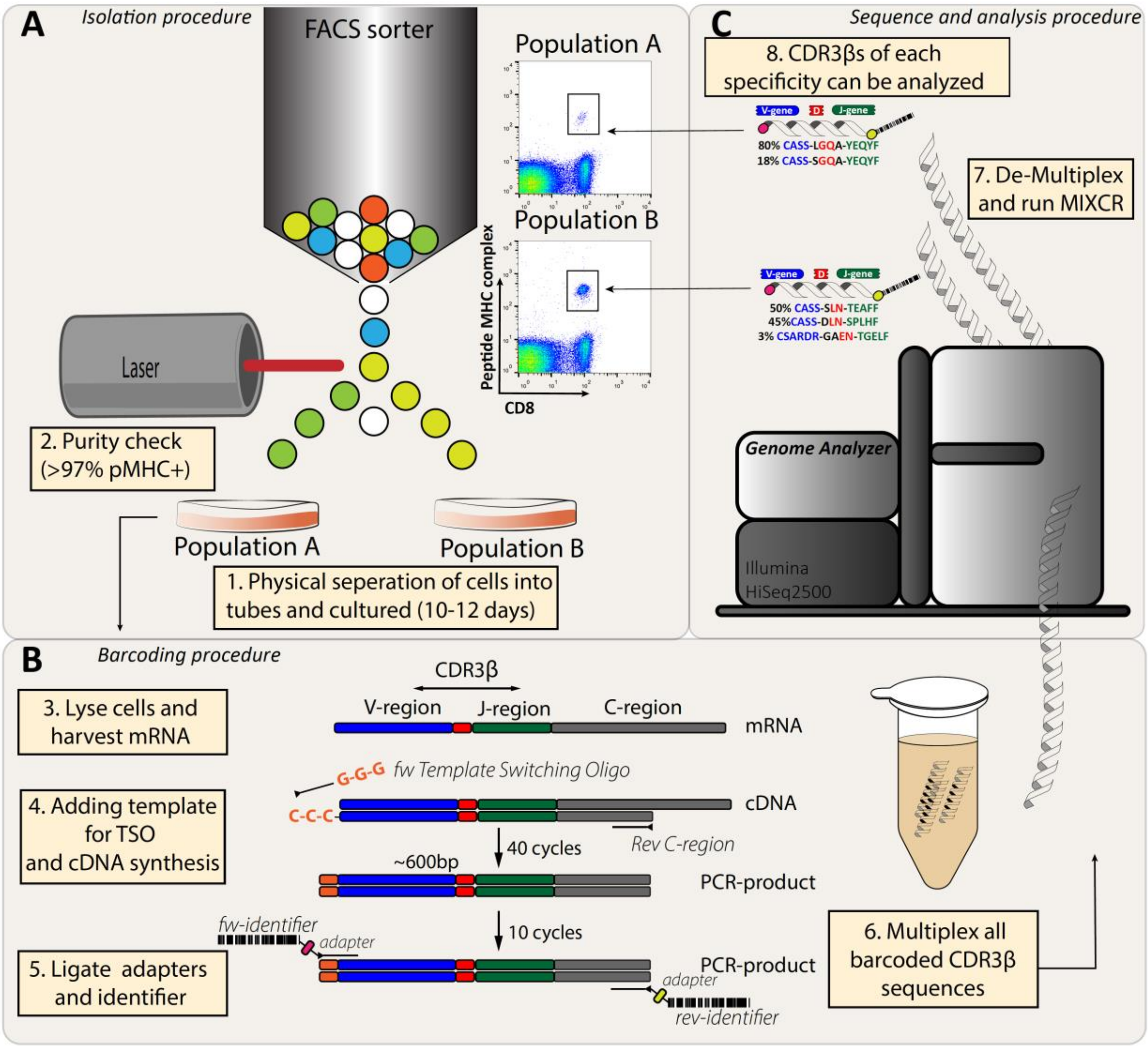

**Supplementary Figure 2: Experimental setup to generate a library of CDR3β-sequences from virus-specific T-cell populations. A**) A total of 190 virus-specific T-cell populations, restricted to HLA-A*01:01, HLA-A*02:01, HLA-B*07:02 or HLA-B*08:01 were isolated using 21 different peptideMHC-tetramers. **B**) Virus-specific T-cell populations were lysed and mRNA was harvested. In the first PCR step, primers specific for the C-region and template switching oligos were added to allow for cDNA synthesis and amplification. A second PCR step was performed with a single primer on each site, which adds unique forward and reverse identifiers (6 basepairs) to each PCR-product for each T-cell population. All 190 PCR-products were multiplexed and high-throughput sequenced. **C**) The library was de-multiplexed based on the unique identifiers. The CDR3β-region was determined using bi-directional readings with MIXCR.

**Supplementary Table 1: Primer sequences.**

| Description | Name | Nucleotide sequence 5'►3' |
|---|---|---|
| cDNA primer TRB constant region reverse transcription | TRB_RT | CACGTGGTCGGGGWAGAAGC |
| cDNA primer SmartSeq2modified template switching oligo | SS2m_TSO | AAGCAGTGGTATCAACGCAGAGTACAT(G)(G){G} |
| PCR primer SmartSeq2modified forward | SS2m_For | GAGTTCAGACGTGTGCTCTTCCGATCTAAGCAGTGGTATC AACGCAGAGTACAT*G |
| PCR primer TRBC1 reverse | TRBC1_rev | CCTACACGACGCTCTTCCGATCTGTGGGAACACCTTGTTCA GGTCCT*C |
| PCR primer TRBC1 reverse | TRBC2_rev | CCTACACGACGCTCTTCCGATCTGTGGGAACACGTTTTTCA GGTCCT*C |
| Barcode primer SS2m region, forward, backbone | BC_R7xx_For | CAAGCAGAAGACGGCATACGAGAT$_{nnnnnn}$GTGACTGGAGTT CAGACGTGTGCTCTTCCGAT*C |
| Barcode primer TRBC region reverse, backbone | BC_R7xx_Rev | AATGATACGGCGACCACCGAGATCTACAC$_{nnnnnn}$ACACTCTT TCCCTACACGACGCTCTTCCGATC*T |

Abbreviations: TRB: T-cell Receptor Beta, SS2m: SmartSeq2Modified, TSO: Template Switching Oligo, TRBC: T-cell Receptor Beta Constant, For: Forward, Rev: Reverse, BC: Beta chain, nnnnnn: Identifier sequence

()=RNA, {}=LNA: Locked Nucleic Acid, *:phosphonothioate-binding

**Supplementary Table 2: Identifier sequences.**

| Identifiers (For) Name | Identifiers (For) Seq | Identifiers (Rev) Name | Identifiers (Rev) Seq |
|---|---|---|---|
| BC_R701 | ATCACG | BC_R725 | ACTGAT |
| BC_R702 | CGATGT | BC_R726 | ATGAGC |
| BC_R703 | TTAGGC | BC_R727 | ATTCCT |
| BC_R704 | TGACCA | BC_R728 | CAAAAG |
| BC_R705 | ACAGTG | BC_R729 | CAACTA |
| BC_R706 | GCCAAT | BC_R730 | CACCGG |
| BC_R707 | CAGATC | BC_R731 | CACGAT |
| BC_R708 | ACTTGA | BC_R732 | CACTCA |
| BC_R709 | GATCAG | BC_R733 | CAGGCG |
| BC_R710 | TAGCTT | BC_R734 | CATGGC |
| BC_R711 | GGCTAC | BC_R735 | CATTTT |
| BC_R712 | CTTGTA | BC_R736 | CCAACA |
| BC_R713 | AGTCAA | BC_R737 | CGGAAT |
| BC_R714 | AGTTCC | BC_R738 | CTAGCT |
| BC_R715 | ATGTCA | BC_R739 | CTATAC |
| BC_R716 | CCGTCC | BC_R740 | CTCAGA |
| BC_R717 | GTAGAG | BC_R741 | GACGAC |
| BC_R718 | GTCCGC | BC_R742 | TAATCG |
| BC_R719 | GTGAAA | BC_R743 | TACAGC |
| BC_R720 | GTGGCC | BC_R744 | TATAAT |
| BC_R721 | GTTTCG | BC_R745 | TCATTC |
| BC_R722 | CGTACG | BC_R746 | TCCCGA |
| BC_R723 | GAGTGG | BC_R747 | TCGAAG |
| BC_R724 | GGTAGC | BC_R748 | TCGGCA |

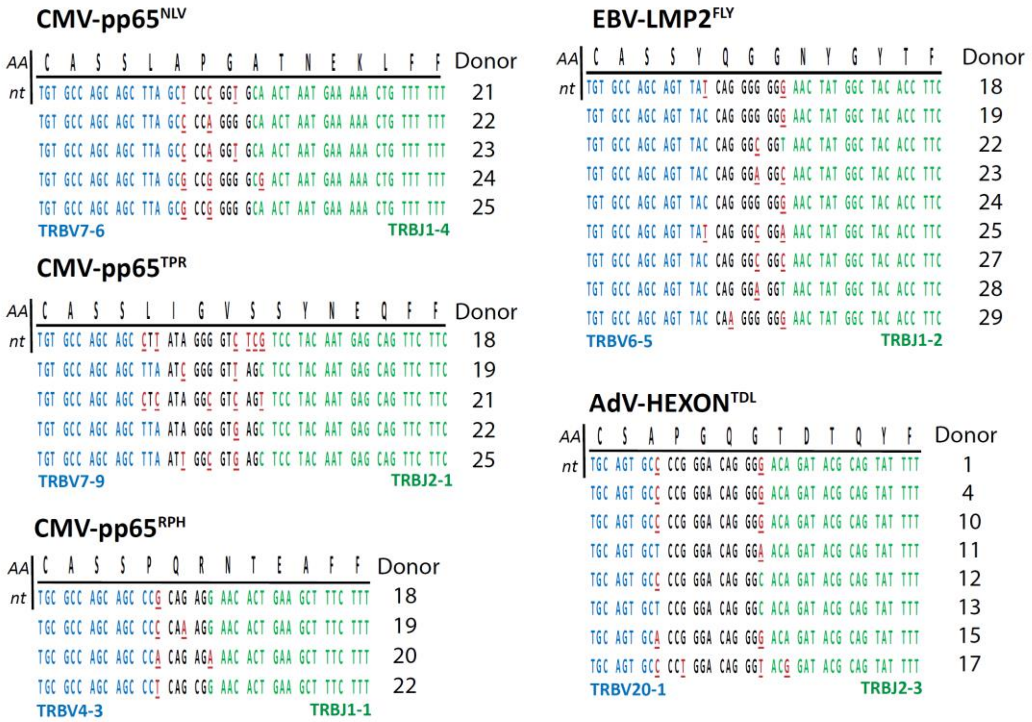

**Germline sequences**

| | | | | | | | |
|---|---|---|---|---|---|---|---|
| **TRBV4-3** | TGCGCCAGCAGCCAAGA | **TRBV7-9** | TGTGCCAGCAGCTTAGC | **TRBJ1-1** | TGAACACTGAAGCTTTCTTT | **TRBJ2-1** | CTCCTACAATGAGCAGTTCTTC |
| **TRBV6-5** | TGTGCCAGCAGTTACTC | **TRBV20-1** | TGCAGTGCTAGAGA | **TRBJ1-2** | CTAACTATGGCTACACCTTC | **TRBJ2-3** | AGCACAGATACGCAGTATTTT |
| **TRBV7-6** | TGTGCCAGCAGCTTAGC | | | **TRBJ1-4** | CAACTAATGAAAAACTGTTTTTT | | |

**Supplementary Figure 3. Identical shared CDR3β amino-acid sequences are found in different individuals with small nucleotide differences as a result of convergent recombination.** The CDR3β nucleotide sequences are shown per donor for 6 identical shared CDR3β amino-acid sequences. Underlined nucleotides in red resemble differences between the different individuals. Nucleotide sequences in blue and green represent perfect alignment with the germline sequences of the TRBV-gene and TRBJ-gene, respectively. The legend represents the germline sequences of (part of) the TRBV and TRBJ genes

**Supplementary table 3: Occurrence and number of CDR3β amino-acid sequences that are shared between individuals.**

| Virus | Antigen | HLA | TRBV | CDR3 | TRBJ | Occurrence (#) |
|---|---|---|---|---|---|---|
| CMV | pp50-VTE | A*01 | TRBV20-1 | CSARLLGGGQSYEQYF | TRBJ2-7 | 2/7 |
| CMV | pp50-VTE | A*01 | TRBV9 | CASSVGQGSSYEQYF | TRBJ2-7 | 2/7 |
| CMV | pp65-YSE | A*01 | TRBV9 | CASSVTGGTDTQYF | TRBJ2-3 | 2/6 |
| CMV | pp65-NLV | A*02 | TRBV7-6 | CASSLAPGATNEKLFF | TRBJ1-4 | 5/8 |
| CMV | pp65-NLV | A*02 | TRBV7-6 | CASSLAPGTTNEKLFF | TRBJ1-4 | 3/8 |
| CMV | pp65-NLV | A*02 | TRBV12-4 | CASSSAYYGYTF | TRBJ1-2 | 2/8 |
| CMV | IE1-VLE | A*02 | TRBV7-3 | CASSLGQGGVETQYF | TRBJ2-5 | 2/6 |
| CMV | IE1-VLE | A*02 | TRBV7-3 | CASSPGQGGVETQYF | TRBJ2-5 | 2/6 |
| CMV | pp65-TPR | B*07 | TRBV7-9 | CASSLIGVSSYNEQFF | TRBJ2-1 | 5/8 |
| CMV | pp65-RPH | B*07 | TRBV4-3 | CASSPQRNTEAFF | TRBJ1-1 | 4/6 |
| CMV | pp65-RPH | B*07 | TRBV4-3 | CASSPSRNTEAFF | TRBJ1-1 | 2/6 |
| CMV | IE1-ELR | B*08 | TRBV27 | CASSSYRTDLNTEAFF | TRBJ1-1 | 2/5 |
| CMV | IE1-QIK | B*08 | TRBV9 | CASSTQVSEPNTGELFF | TRBJ2-2 | 2/6 |
| CMV | IE1-QIK | B*08 | TRBV9 | CASSVQRQTANTGELFF | TRBJ2-2 | 2/6 |
| CMV | IE1-QIK | B*08 | TRBV12-5 | CASGPRAGAYNEQFF | TRBJ2-1 | 2/6 |
| CMV | IE1-QIK | B*08 | TRBV2 | CASSGTGRLTMNTEAFF | TRBJ1-1 | 2/6 |
| CMV | IE1-QIK | B*08 | TRBV21-1 | CASSKVAARVP-TLKLS | TRBJ1-1 | 2/6 |
| CMV | IE1-QIK | B*08 | TRBV7-9 | CASSLTLAGNQPQHF | TRBJ1-5 | 2/6 |
| EBV | LMP2-FLY | A*02 | TRBV6-5 | CASSYQGGNYGYTF | TRBJ1-2 | 9/11 |
| EBV | LMP2-FLY | A*02 | TRBV6-5 | CASSRQGGNYGYTF | TRBJ1-2 | 7/11 |
| EBV | LMP2-FLY | A*02 | TRBV6-5 | CASSLQGGNYGYTF | TRBJ1-2 | 5/11 |
| EBV | LMP2-FLY | A*02 | TRBV6-5 | CASSGQGGNYGYTF | TRBJ1-2 | 4/11 |
| EBV | LMP2-FLY | A*02 | TRBV6-5 | CASSKQGGGYGYTF | TRBJ1-2 | 3/11 |
| EBV | LMP2-FLY | A*02 | TRBV6-5 | CASSPQGGGYGYTF | TRBJ1-2 | 3/11 |
| EBV | LMP2-FLY | A*02 | TRBV6-5 | CASSRQGGTYGYTF | TRBJ1-2 | 3/11 |
| EBV | LMP2-FLY | A*02 | TRBV6-5 | CASSSQGGNYGYTF | TRBJ1-2 | 3/11 |
| EBV | LMP2-FLY | A*02 | TRBV6-5 | CASSYSGGYYGYTF | TRBJ1-2 | 2/11 |
| EBV | LMP2-FLY | A*02 | TRBV6-5 | CASSDQGGGYGYTF | TRBJ1-2 | 2/11 |
| EBV | LMP2-FLY | A*02 | TRBV6-5 | CASSFQGGNYGYTF | TRBJ1-2 | 2/11 |
| EBV | LMP2-FLY | A*02 | TRBV6-5 | CASSPLGGAEGYTF | TRBJ1-2 | 2/11 |
| EBV | BRLF1-YVL | A*02 | TRBV10-1 | CASSAGPDTQYF | TRBJ2-3 | 2/12 |
| EBV | BRLF1-YVL | A*02 | TRBV24-1 | CATSDYGEDTQYF | TRBJ2-3 | 2/12 |
| EBV | BRLF1-YVL | A*02 | TRBV25-1 | CASSEWTTDTQYF | TRBJ2-3 | 2/12 |
| EBV | BRLF1-YVL | A*02 | TRBV28 | CASSKIMNTEAFF | TRBJ1-1 | 2/12 |
| EBV | BRLF1-YVL | A*02 | TRBV6-5 | CASSQLLGSNQPQHF | TRBJ1-5 | 2/12 |
| EBV | EBNA3A-RPP | B*07 | TRBV4-1 | CASSQDRLTGGYTF | TRBJ1-2 | 4/11 |
| EBV | EBNA3A-RPP | B*07 | TRBV4-1 | CASSQDRLTGTQYF | TRBJ2-5 | 2/11 |
| EBV | EBNA3A-RPP | B*07 | TRBV4-1 | CASSQEAFNYEQYF | TRBJ2-7 | 2/11 |
| EBV | BZLF1-RAK | B*08 | TRBV27 | CASSSLNTEAFF | TRBJ1-1 | 8/17 |
| EBV | BZLF1-RAK | B*08 | TRBV27 | CASSPLTDTQYF | TRBJ2-3 | 3/17 |
| EBV | BZLF1-RAK | B*08 | TRBV20-1 | CSARDRGAENTGELFF | TRBJ2-2 | 3/17 |
| EBV | BZLF1-RAK | B*08 | TRBV20-1 | CSARDRGGENTGELFF | TRBJ2-2 | 3/17 |
| EBV | BZLF1-RAK | B*08 | TRBV29-1 | CSVGSGEGYEQYF | TRBJ2-7 | 3/17 |
| EBV | BZLF1-RAK | B*08 | TRBV4-1 | CASSPGQGEGYEQYF | TRBJ2-7 | 3/17 |
| EBV | BZLF1-RAK | B*08 | TRBV7-9 | CASSPTGAGNQPQHF | TRBJ1-5 | 3/17 |
| EBV | BZLF1-RAK | B*08 | TRBV27 | CASSNLNTEAFF | TRBJ1-1 | 2/17 |
| EBV | BZLF1-RAK | B*08 | TRBV27 | CASSDLNSPLHF | TRBJ1-6 | 2/17 |
| EBV | BZLF1-RAK | B*08 | TRBV27 | CASSSLNSPLHF | TRBJ1-6 | 2/17 |
| EBV | BZLF1-RAK | B*08 | TRBV7-2 | CASSLVLLGNSPLHF | TRBJ1-6 | 2/17 |
| EBV | BZLF1-RAK | B*08 | TRBV4-1 | CASSRLAGDTDTQYF | TRBJ2-3 | 2/17 |
| EBV | BZLF1-RAK | B*08 | TRBV6-1 | CASTGTASTDTQYF | TRBJ2-3 | 2/17 |
| EBV | BZLF1-RAK | B*08 | TRBV20-1 | CSARDRGSENTGELFF | TRBJ2-2 | 2/17 |
| EBV | BZLF1-RAK | B*08 | TRBV20-1 | CSARDRGTENTGELFF | TRBJ2-2 | 2/17 |
| EBV | BZLF1-RAK | B*08 | TRBV7-3 | CASSSHSGINTGELFF | TRBJ2-2 | 2/17 |
| EBV | BZLF1-RAK | B*08 | TRBV10-3 | CATGLAGSTDTQYF | TRBJ2-3 | 2/17 |
| EBV | BZLF1-RAK | B*08 | TRBV4-1 | CASSPGTGEGYEQYF | TRBJ2-7 | 2/17 |
| EBV | BZLF1-RAK | B*08 | TRBV7-2 | CASSPGTGEGYEQYF | TRBJ2-7 | 2/17 |
| EBV | BZLF1-RAK | B*08 | TRBV7-2 | CASSYHGSYEQYF | TRBJ2-7 | 2/17 |
| EBV | BZLF1-RAK | B*08 | TRBV7-6 | CASSLAGEGYEQYF | TRBJ2-7 | 2/17 |
| EBV | BZLF1-RAK | B*08 | TRBV7-9 | CASSSTGAGNQPQHF | TRBJ1-5 | 2/17 |
| EBV | BZLF1-RAK | B*08 | TRBV7-9 | CASSSTGSGDQPQHF | TRBJ1-5 | 2/17 |
| EBV | BZLF1-RAK | B*08 | TRBV7-3 | CASSLIASGGYNEQFF | TRBJ2-1 | 2/17 |
| EBV | EBNA3A-FLR | B*08 | TRBV7-8 | CASSLGQAYEQYF | TRBJ2-7 | 4/13 |
| EBV | EBNA3A-FLR | B*08 | TRBV7-8 | CASSSGQAYEQYF | TRBJ2-7 | 4/13 |
| EBV | EBNA3A-FLR | B*08 | TRBV7-8 | CASSTGQAYEQYF | TRBJ2-7 | 3/13 |

| | | | | | | |
|---|---|---|---|---|---|---|
| EBV | LMP2-FLY | A*02 | TRBV6-5 | CASSPQGGNYGYTF | TRBJ1-2 | 2/11 |
| EBV | LMP2-FLY | A*02 | TRBV6-5 | CASSPQGGRDGYTF | TRBJ1-2 | 2/11 |
| EBV | LMP2-FLY | A*02 | TRBV6-5 | CASSRQGGSYGYTF | TRBJ1-2 | 2/11 |
| EBV | LMP2-FLY | A*02 | TRBV6-5 | CASSSQGGSNYGYTF | TRBJ1-2 | 2/11 |
| EBV | LMP2-FLY | A*02 | TRBV6-5 | CASSSQGGSYGYTF | TRBJ1-2 | 2/11 |
| EBV | LMP2-FLY | A*02 | TRBV6-5 | CASSYEGGYYGYTF | TRBJ1-2 | 2/11 |
| EBV | LMP2-FLY | A*02 | TRBV6-5 | CASSYQGGSYGYTF | TRBJ1-2 | 2/11 |
| EBV | LMP2-FLY | A*02 | TRBV6-5 | CASNPQGGGGGYTF | TRBJ1-2 | 2/11 |
| EBV | LMP2-FLY | A*02 | TRBV6-5 | CASNPQGGGNGYTF | TRBJ1-2 | 2/11 |
| EBV | LMP2-FLY | A*02 | TRBV6-5 | CASSYQGGNEQFF | TRBJ2-1 | 3/11 |
| EBV | LMP2-FLY | A*02 | TRBV6-5 | CASSLQGGNEQFF | TRBJ2-1 | 2/11 |
| EBV | LMP2-FLY | A*02 | TRBV6-5 | CASTLQGGNEQFF | TRBJ2-1 | 2/11 |
| EBV | LMP2-CLG | A*02 | TRBV10-2 | CASSEDGMNTEAFF | TRBJ1-1 | 3/10 |
| EBV | LMP2-CLG | A*02 | TRBV10-2 | CASSSDGMNTEAFF | TRBJ1-1 | 2/10 |
| EBV | LMP2-CLG | A*02 | TRBV10-2 | CASSGDGMNTEAFF | TRBJ1-1 | 2/10 |
| EBV | LMP2-CLG | A*02 | TRBV10-2 | CASSQDGMNTEAFF | TRBJ1-1 | 2/10 |
| EBV | LMP2-CLG | A*02 | TRBV5-1 | CASSLEGQASSYEQYF | TRBJ2-7 | 3/10 |
| EBV | EBNA3C-LLD | A*02 | TRBV19 | CASSIALASEQYF | TRBJ2-7 | 2/7 |
| EBV | BMLF1-GLC | A*02 | TRBV29-1 | CSVGTGGTNEKLFF | TRBJ1-4 | 6/10 |
| EBV | BMLF1-GLC | A*02 | TRBV20-1 | CSARDRVGNTIYF | TRBJ1-3 | 5/10 |
| EBV | BMLF1-GLC | A*02 | TRBV20-1 | CSARDGTGNGYTF | TRBJ1-2 | 3/10 |
| EBV | BMLF1-GLC | A*02 | TRBV20-1 | CSARDRTGNGYTF | TRBJ1-2 | 3/10 |
| EBV | BMLF1-GLC | A*02 | TRBV29-1 | CSVGAGGTNEKLFF | TRBJ1-4 | 3/10 |
| EBV | BMLF1-GLC | A*02 | TRBV14 | CASSQSPGGTQYF | TRBJ2-3 | 2/10 |
| EBV | BMLF1-GLC | A*02 | TRBV20-1 | CSARVGVGNTIYF | TRBJ1-3 | 2/10 |
| EBV | BMLF1-GLC | A*02 | TRBV29-1 | CSAGSGGTNEKLFF | TRBJ1-4 | 2/10 |
| EBV | BMLF1-GLC | A*02 | TRBV29-1 | CSVGSGGTNEKLFF | TRBJ1-4 | 2/10 |
| EBV | BRLF1-YVL | A*02 | TRBV20-1 | CSAIGGSYNEQFF | TRBJ2-1 | 3/12 |
| EBV | BRLF1-YVL | A*02 | TRBV20-1 | CSAPVPPYNEQFF | TRBJ2-1 | 2/12 |
| EBV | BRLF1-YVL | A*02 | TRBV20-1 | CSARGTEFYEQYF | TRBJ2-7 | 2/12 |
| EBV | BRLF1-YVL | A*02 | TRBV28 | CASSLFSNEQFF | TRBJ2-1 | 2/12 |

| | | | | | | |
|---|---|---|---|---|---|---|
| EBV | EBNA3A-FLR | B*08 | TRBV4-3 | CASSHGLAGILETQYF | TRBJ2-5 | 2/13 |
| EBV | EBNA3A-FLR | B*08 | TRBV4-3 | CASSPTSGVAGELFF | TRBJ2-2 | 2/13 |
| EBV | EBNA3A-FLR | B*08 | TRBV4-1 | CASSQGLAVSSYEQYF | TRBJ2-7 | 2/13 |
| EBV | EBNA3A-FLR | B*08 | TRBV7-9 | CASSWGPEQFF | TRBJ2-1 | 2/13 |
| EBV | EBNA3A-QAK | B*08 | TRBV18 | CAASRGCEPKTFST | TRBJ2-4 | 5/18 |
| EBV | EBNA3A-QAK | B*08 | TRBV28 | CASSNLGVTELNTGELFF | TRBJ2-2 | 3/18 |
| EBV | EBNA3A-QAK | B*08 | TRBV5-1 | CASSLELAVYNEQFF | TRBJ2-1 | 3/18 |
| EBV | EBNA3A-QAK | B*08 | TRBV5-1 | CASSLETATEAFF | TRBJ1-1 | 3/18 |
| EBV | EBNA3A-QAK | B*08 | TRBV5-1 | CASSLETGGYGYTF | TRBJ1-2 | 3/18 |
| EBV | EBNA3A-QAK | B*08 | TRBV27 | CASSLYRDNQPQHF | TRBJ1-5 | 2/18 |
| EBV | EBNA3A-QAK | B*08 | TRBV27 | CASSPDRWETQYF | TRBJ2-5 | 2/18 |
| EBV | EBNA3A-QAK | B*08 | TRBV28 | CASSALSGLAGPGELFF | TRBJ2-2 | 2/18 |
| EBV | EBNA3A-QAK | B*08 | TRBV28 | CASSKQGAPGHTGELFF | TRBJ2-2 | 2/18 |
| EBV | EBNA3A-QAK | B*08 | TRBV28 | CASSLLGARGLNEKLFF | TRBJ1-4 | 2/18 |
| EBV | EBNA3A-QAK | B*08 | TRBV28 | CASSLLGTGGLSEKLFF | TRBJ1-4 | 2/18 |
| EBV | EBNA3A-QAK | B*08 | TRBV28 | CASSQQGARSLSEKLFF | TRBJ1-4 | 2/18 |
| EBV | EBNA3A-QAK | B*08 | TRBV4-2 | CASSQDAGDRLAGVTGELFF | TRBJ2-2 | 2/18 |
| EBV | EBNA3A-QAK | B*08 | TRBV5-1 | CASSLETGDTQYF | TRBJ2-3 | 2/18 |
| EBV | EBNA3A-QAK | B*08 | TRBV6-3 | CASSLDPPGQSIRVNTGELFF | TRBJ2-2 | 2/18 |
| AdV | HEXON-TDL | A*01 | TRBV20-1 | CSAPGQGTDTQYF | TRBJ2-3 | 8/12 |
| AdV | HEXON-TDL | A*01 | TRBV20-1 | CSAPGQGTTEAFF | TRBJ1-1 | 4/12 |
| AdV | HEXON-TDL | A*01 | TRBV20-1 | CSAPGQGTYEQYF | TRBJ2-7 | 3/12 |
| AdV | HEXON-TDL | A*01 | TRBV20-1 | CSAPGQGSTEAFF | TRBJ1-1 | 3/12 |
| AdV | HEXON-TDL | A*01 | TRBV20-1 | CSAPGQGEETQYF | TRBJ2-5 | 2/12 |
| AdV | HEXON-TDL | A*01 | TRBV20-1 | CSAPGQGENTQYF | TRBJ2-3 | 2/12 |
| AdV | HEXON-TDL | A*01 | TRBV5-1 | CASNDYDNEQFF | TRBJ2-1 | 2/12 |
| AdV | HEXON-TDL | A*01 | TRBV5-1 | CASNLADDEQFF | TRBJ2-1 | 2/12 |
| AdV | HEXON-TDL | A*01 | TRBV10-3 | CATQTGGSNQPQHF | TRBJ1-5 | 2/12 |
| AdV | HEXON-TDL | A*01 | TRBV4-1 | CASSQVVGQAHSPLHF | TRBJ1-6 | 2/12 |
| AdV | HEXON-TDL | A*01 | TRBV6-6 | CASSYPGNNSPLHF | TRBJ1-6 | 2/12 |
| AdV | HEXON-TDL | A*01 | TRBV20-1 | CSAR-ASVATSST | TRBJ2-7 | 2/12 |
| AdV | HEXON-TDL | A*01 | TRBV19 | CATSSAAQETQYF | TRBJ2-5 | 2/12 |
| AdV | HEXON-KPY | B*07 | TRBV10-3 | CAINPGTAYGYTF | TRBJ1-2 | 2/8 |
| AdV | HEXON-KPY | B*07 | TRBV18 | CASSPGTPEQFF | TRBJ2-1 | 2/8 |

A total of 131 different shared identical CDR3β amino-acid sequences are shown. The number of T-cell populations that contain the shared identical CDR3β amino-acid sequences are shown per total number of T-cell populations of that respective specificity (#), reflecting the occurrence among donors.

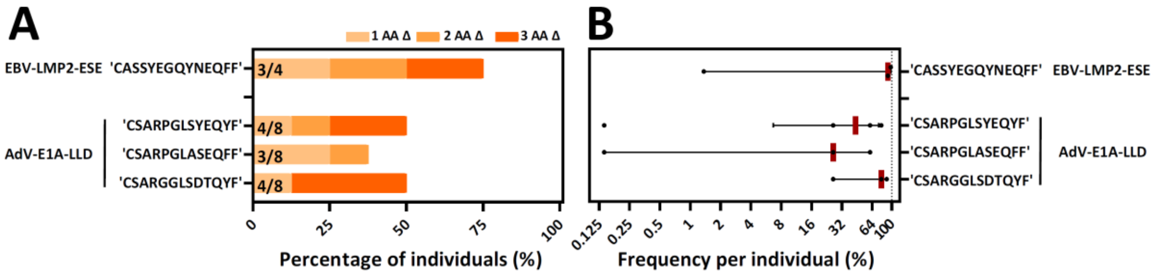

**Supplementary figure 4. Highly similar CDR3β amino-acid sequences in specific T-cell populations that did not contain an identical shared CDR3β amino-acid sequence**. **A)** For two specificities no identical shared CDR3β amino-acid sequences were found.. Individuals did contain highly similar sequences, and these were stacked with 1, 2 or 3 amino-acid differences. The occurrence, shown as percentages among healthy donors, is shown per CDR3β amino-acid sequence. The total number of different T-cell populations (different donors) for each specificity/CDR3β amino-acid sequence is shown at the inner-side of the y-axis. **B)** Shown is the sum of frequencies of the identical and highly similar (1,2 and 3 amino-acid differences) CDR3β amino-acid sequences per individual. Each dot is one individual, and the red-lines represents the medians with interquartile ranges

AA: amino-acids, nt: nucleotides, Δ: difference(s)
